# Genetic and chemotherapeutic causes of germline hypermutation

**DOI:** 10.1101/2021.06.01.446180

**Authors:** Joanna Kaplanis, Benjamin Ide, Rashesh Sanghvi, Matthew Neville, Petr Danecek, Tim Coorens, Elena Prigmore, Patrick Short, Giuseppe Gallone, Jeremy McRae, Chris Odhams, Loukas Moutsianas, Genomics England Research Consortium, Jenny Carmichael, Angela Barnicoat, Helen Firth, Patrick O’Brien, Raheleh Rahbari, Matthew Hurles

## Abstract

Mutation in the germline is the source of all evolutionary genetic variation and a cause of genetic disease. Previous studies have shown parental age to be the primary determinant of the number of new germline mutations seen in an individual’s genome. Here we analysed the genome-wide sequences of 21,879 families with rare genetic diseases and identified 12 hypermutated individuals with between two and seven times more *de novo* single nucleotide variants (dnSNVs) than expected. In most of these families (8/12) the excess mutations could be attributed to the father. We determined that two of these families had genetic drivers of germline hypermutation, with the fathers carrying damaging genetic variation in known DNA repair genes, causing distinctive mutational signatures. For five families, by analysing clinical records and mutational signatures, we determined that paternal exposure to chemotherapeutic agents prior to conception was a key driver of hypermutation. Our results suggest that the germline is well protected from mutagenic effects, hypermutation is rare and relatively modest in degree and that most hypermutated individuals will not have a genetic disease.

## Introduction

Germline mutagenesis is the source of all genetic variation which drives evolution and generates disease-causing variants. The average number of *de novo* mutations (DNMs) generating single nucleotide variants (SNVs) is estimated to be 60-70 per human genome per generation, but little is known about germline hypermutated individuals with unusually large numbers of DNMs^1–3^. The human germline mutation rate is not a constant, but varies between individuals, families and populations and has evolved over time just like any other phenotype^45–8^. Parental age explains a large proportion of variance for single nucleotide variants (SNVs), indels and short tandem repeats (STRs)^3,9,10^ It has been estimated that there is an increase of ~2 DNMs for every additional year in father’s age and a more subtle increase of ~0.5 DNMs for every additional year in mother’s age^3,11^. Subtle differences have also been observed between the maternal and paternal mutational spectra and may be indicative of different mutagenic processes^12–15^. Different mutational mechanisms can leave distinct mutational patterns. These combinations of mutation types can be decomposed from mutational spectra into ‘mutational signatures’^16,17^. There are currently >100 somatic mutational signatures that have been identified across a wide variety of cancers of which half have been attributed to endogenous mutagenic processes or specific mutagens^18,19^. The majority of germline mutation can be explained by two of these signatures, termed signature 1 (SBS1), likely due to deamination of 5-methylcytosine^20^, and signature 5 (SBS5), thought to be a pervasive and relatively clock-like endogenous process. Both signatures are ubiquitous among normal and cancer cell types^21,22^ and have been reported previously in trio-studies^13^. The impact of environmental mutagens has been well established in the soma but is not as well understood in the germline^23,24^. Environmental exposures in parents, such as ionising radiation, can influence the number of mutations transmitted to offspring^25–27^. Individual mutation rates can also be influenced by genetic background. With regards to somatic mutation, thousands of inherited germline variants have been shown to increase cancer risk^28–30^. Many of these variants are in genes encoding components of DNA repair pathways which, when impaired, lead to an increased number of somatic mutations. However it is not known whether variants in known somatic mutator genes can influence germline mutation rates. There are a handful of examples where genetic background has been shown to impact the germline mutation rate of STRs, minisatellites and translocations, often in cis, rather than genome-wide^31–34,35^.

An elevated germline mutation rate can have a significant impact on the health of subsequent generations. Increasing germline mutation rate results in an increased risk of offspring being born with a genetic disorder caused by a DNM^36^. Long-term effects of mutation rate differences as a result of mutation accumulation have been demonstrated in mice to have effects on reproduction and survival rates and there may be a similar impact in humans^37,38^.

While we have started to explain the general properties of germline mutations, little is known about rare outliers with extreme mutation rates. *De novo* mutations are a substantial cause of rare genetic disorders and cohorts of patients with such disorders are enriched for DNMs overall and are more likely to include germline hypermutated individuals^11,39^. To this end we sought to identify germline hypermutated individuals in ~20,000 sequenced parent offspring trios from two rare disease cohorts. We identified genetic or environmental causes of this hypermutation and estimated how much variation in germline mutation rate this may explain.

## Results

### Identifying germline hypermutated individuals in rare disease cohorts

We sought to identify germline hypermutated individuals in two separate cohorts: 7,930 exome-sequenced parent offspring trios from the Deciphering Developmental Disorders (DDD) Study and 13,949 whole-genome sequenced parent offspring trios in the rare disease arm of the 100,000 Genome Project (100kGP). We selected nine trios from the DDD study with the largest number of exonic DNMs in the offspring, given their parental ages, which were subsequently whole genome sequenced at >30X coverage to characterise DNMs genome-wide. In the 100kGP cohort, we performed extensive filtering of the DNMs which resulted in a total of 903,525 *de novo* SNVs (dnSNVs) and 72,110 *de novo* indels (dnIndels). The median number of DNMs per individual was 62 for dnSNVs and 5 for dnIndels (median paternal and maternal ages of 33 and 30) (Supplemental Figure 1).

Parental age explains the majority of variance in numbers of germline mutations observed in offspring and is important to control for when examining additional sources of variation^3^. We observed an increase in total number of dnSNVs of 1.28 dnSNVs/year of paternal age (CI:1.24-1.32, p<10^-300^, Negative binomial regression) and an increase of 0.35 dnSNVs/year of maternal age (CI: 0.30-0.39, p = 3.0 ×10^-49^, Negative Binomial regression) (Figure 1 a). We were able to phase 241,063 dnSNVs and found that 77% of phased DNMs were paternal in origin, which agrees with previous estimates^12–14^. Estimates of the parental age effect in the phased mutations were not significantly different to the unphased results: 1.23 paternal dnSNVs/year of paternal age (CI: 1.14-1.32, p =1.6×10^-158^) and 0.38 maternal dnSNVs/year of maternal age (CI: 0.35,0.41, p = 6.6×10^-120^) (Supplemental Figure 2b). Paternal and maternal age were also significantly associated with the number of dnIndels: an increase of 0.071 dnIndels/year of paternal age (CI: 0.062-0.080, p = 8.3×10^-56^, Supplemental Figure 2a) and a smaller increase of 0.019 dnIndels/year of maternal age (CI: 0.0085-0.029 p = 3.4 ×10^-4^, Supplemental Figure 2a). The ratio of paternal to maternal mutation increases for SNVs and indels were very similar, 3.7 for SNVs and 3.8 for indels. The proportion of *de novo* mutations that phased paternally increased by 0.0017 for every year of paternal age (p = 3.37 ×10^-38^, Binomial regression, Supplemental Figure 3). However, the effect size is small and the proportion of DNMs that phase paternally in the youngest fathers is ~0.75 and so the paternal age effect alone does not fully explain the strong paternal bias^14^. We compared the mutational spectra of the phased DNMs and found that maternally derived DNMs have a significantly higher proportion of C>T mutations (0.27 maternal vs 0.22 paternal, p = 3.24×10^-80^, Binomial test), while paternally derived DNMs have a significantly higher proportion of C>A, T>G and T>C mutations (C>A: 0.08 maternal vs 0.10 paternal, p = 4.6×10^-23^; T>G 0.06 vs 0.7, p = 6.8×10^-28^; T>C 0.25 vs 0.26, p = 1.6×10^-5^; Binomial test, Supplemental Figure 4a). These mostly agree with previous studies although the difference in T>C mutations was not previously significant^12^. The majority of both paternal and maternal mutations could be explained by Signature 1 and 5, with a slightly higher contribution of signature 1 in paternal mutations (0.16 paternal vs 0.15 maternal, chi-squared test p = 2.0 × 10^-5^,, Supplemental Figure 4b).

**Figure 1:**
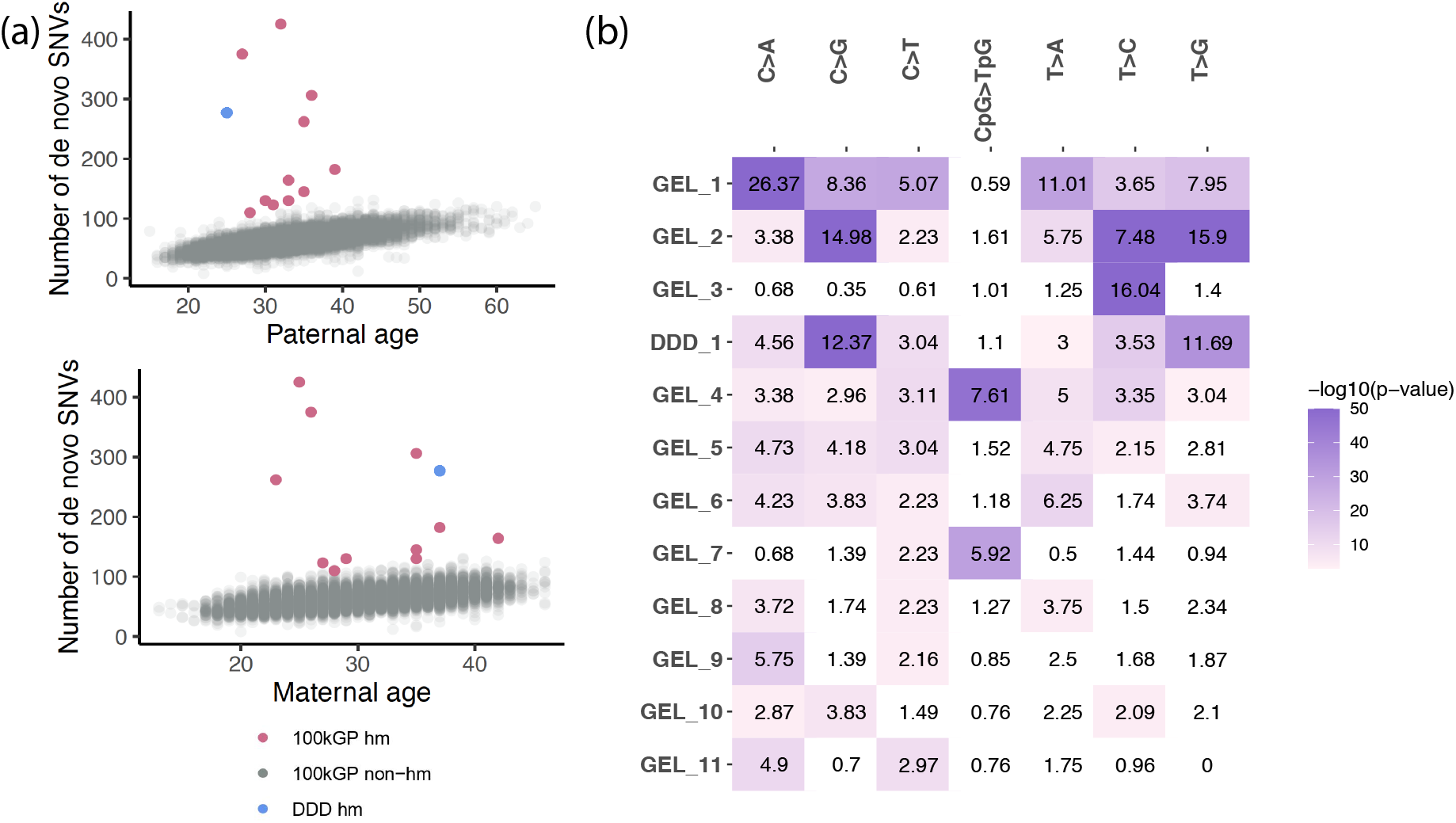
Identification of germline hypermutated individuals. (a) Paternal and maternal age vs number of dnSNVs, 100kGP hypermutated individuals are highlighted in pink and DDD hypermutated individual is highlighted in blue (b) Enrichment (observed/expected) of mutation type for hypermutated individuals. Sample names on the y-axis, mutation type on the x-axis. The enrichment is colored by the -log10(enrichment p-value) which was calculated using a Poisson test comparing the average number of mutations in each type across all individuals in the 100kGP cohort. White coloring indicates no statistically significant enrichment (p-value <0.05/12*7).

We identified 12 germline hypermutated individuals after accounting for parental age (see Methods): 11 from the 100kGP cohort and 1 from the DDD cohort (Figure 1a, Table 1). The number of DNMs genome-wide for each of the 12 hypermutated individuals ranged from 110-425 dnSNVs, which corresponds to a fold increase of 1.7-6.5 compared to the median number of dnSNVs per individual across the 100kGP cohort. Two of these individuals also had a significantly increased number of dnIndels (Table 1). The mutational spectra across these hypermutated individuals varied dramatically (Figure 1b, Supplemental Figure 5, Supplemental Table 1) and after extracting mutational signatures we found that while many of the mutations mapped onto several known somatic signatures (from the Catalogue of Somatic Mutations in Cancer (COSMIC)^40^), a novel mutational signature, termed SBSHYP, was also extracted (Figure 2a,b, Supplemental Table 2). In addition to mutational spectra, we evaluated the parental phase, transcriptional strand bias (Supplemental Figure 6) and the distribution of the variant allele fraction (VAF) for these mutations (Supplemental Figure 7). Upon examining these properties, we identified three potential sources of germline hypermutation: paternal defects in DNA repair genes, paternal exposure to chemotherapeutics and post-zygotic mutational factors.

**Figure 2:**
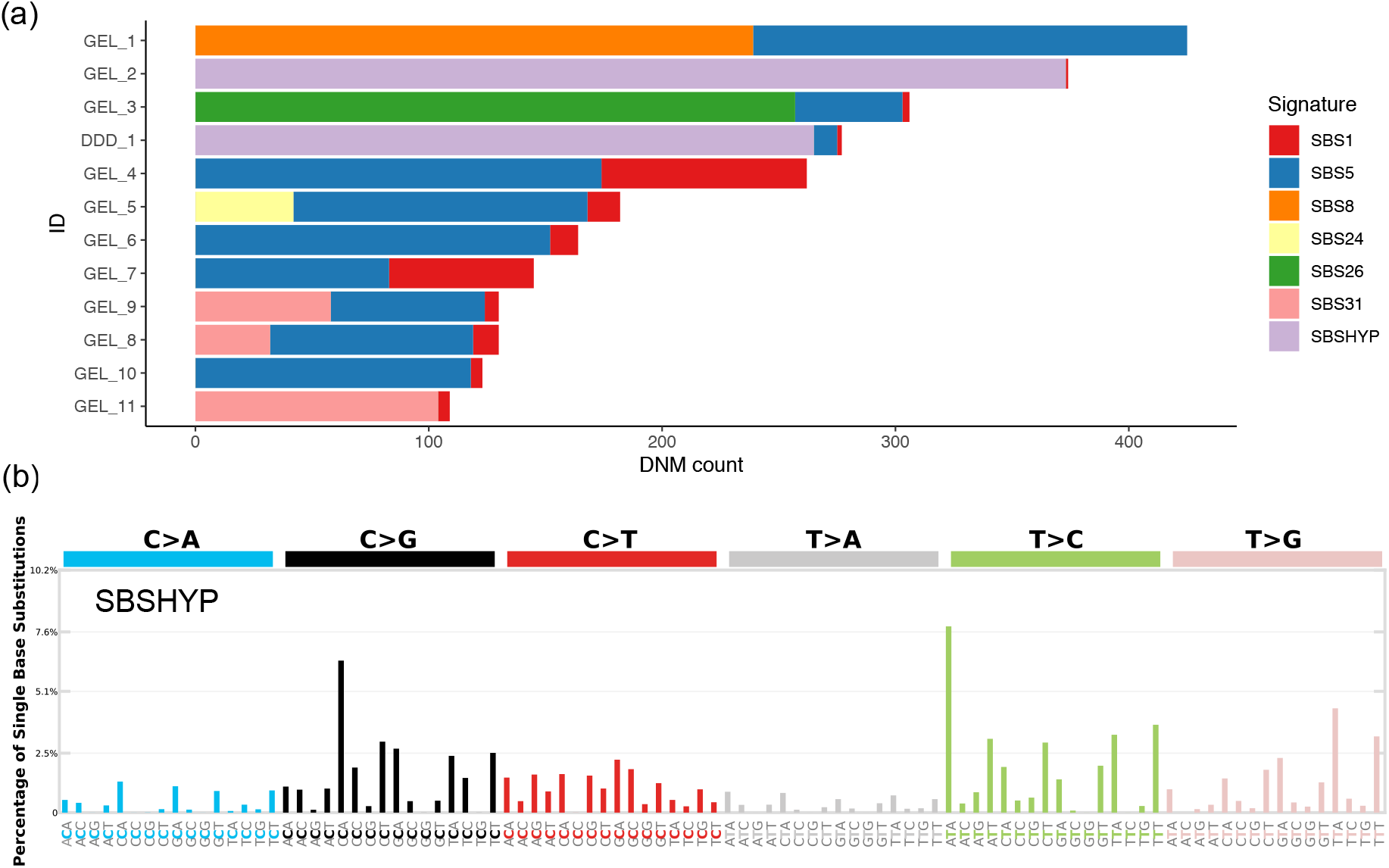
Mutational signatures in 12 germline hypermutated individuals. (a) Contributions of mutational signatures extracted using SigProfiler and decomposed on to known somatic mutational signatures as well as the novel signature (SBSHYP) identified in both DDD_1 and GEL_2. Summary of signatures: SBS1 and SBS5 are known germline signatures; SBS8 associated with TC-NER; SBS26 associated with defective MMR; SBS31 associated with platinum drug treatment; SBS24 is associated with aflatoxin exposure. (b) Trinucleotide context mutational profile of novel extracted mutational signature SBSHYP

**Table 1:** Properties and possible hypermutation sources for 12 germline hypermutated individuals. Eleven individuals were identified in 100kGP as having a significantly large number of dnSNVs (GEL_1-GEL_11) and one individual identified in the DDD study (DDD_1). The DNM counts are for autosomal DNMs. Child age refers to age when sample was taken. Paternal and maternal age refer to age at child’s birth. All ages are given as 5 year ranges for 100kGP individuals and the exact age for the DDD individuals. SNV and indel p-value is from testing the number of dnSNVs and dnIndels compared to what we would expect after accounting for parental age. TS bias: transcriptional strand bias p-value for dnSNVS. Phase (P,M): the number of dnSNVs that phase paternally (P) and maternally (M) with significance indicator for how different this ratio is compared to the observed proportion across all DNMs that phase paternally in 100kGP (0.77) using a Binomial test (*p<0.1, **p<0.01,***p<0.001). We have detailed the parental cancer and chemotherapy drugs received when relevant. Treatments abbreviations: BEP (Bleomycin, etoposide and platinum), ABVD (Bleomycin-Dacarbazine-Doxorubicin-Vinblastine) and IVE (Iphosphamide, epirubicin and etoposide).

| ID | Number of dnSNVs/<br>dnIndels | Child<br>age | Paternal<br>age | Maternal<br>age | SNV<br>p-value | Indel<br>p-value | TS bias | Phase<br>(P,M) | Potential source of<br>hypermutation |
| --- | --- | --- | --- | --- | --- | --- | --- | --- | --- |
| GEL_1 | 425/16 | 5-10 | 30-35 | 20-25 | 4.2e-90 | 9.4e-05 | 2.1e-40 | 129,1*** | Paternal DNA repair defect; homozygous stop-gain *XPC* variant |
| GEL_2 | 375/5 | 10-15 | 25-30 | 25-30 | 2.3e-83 | 0.43 | 0.22 | 106,7*** | Paternal chemotherapy; Nephrotic syndrome: Cyclophosphamide, Chlorambucil |
| GEL_3 | 306/4 | 0-5 | 35-40 | 30-35 | 2.5e-44 | 0.73 | 0.86 | 87,5*** | Paternal DNA repair defect; homozygous missense *MPG* variant |
| DDD_1 | 277/6 | 6 | 25 | 37 | NA | NA | 3.3e-03 | 72,4*** | Paternal chemotherapy; Hodgkins Lymphoma: ABVD, IVE |
| GEL_4 | 262/12 | 10-15 | 30-35 | 20-25 | 1.7e-37 | 0.007 | 0.070 | 36,32*** | Post-zygotic hypermutation |
| GEL_5 | 182/8 | 0-5 | 35-40 | 35-40 | 8.4e-14 | 0.19 | 0.15 | 63,4*** | Paternal chemotherapy; SLE: drugs unknown |
| GEL_6 | 164/7 | 0-5 | 30-35 | 40-45 | 9.8e-13 | 0.25 | 0.066 | 38,3* | Unknown |
| GEL_7 | 145/9 | 0-5 | 30-35 | 30-35 | 2.4e-09 | 0.08 | 0.02 | 24,16* | Post-zygotic hypermutation |
| GEL_8 | 130/6 | 20-25 | 25-30 | 25-30 | 2.1e-09 | 0.31 | 1.00 | 31,11 | Paternal chemotherapy; Testicular cancer: drugs unknown |
| GEL_9 | 130/5 | 5-10 | 30-35 | 30-35 | 1.2e-07 | 0.53 | 0.016 | 46,2** | Paternal chemotherapy; Testicular cancer: BEP |
| GEL_10 | 123/5 | 10-15 | 30-35 | 25-30 | 5.3e-08 | 0.48 | 0.082 | 38,0*** | Unknown |
| GEL_11 | 110/5 | 10-15 | 25-30 | 25-30 | 8.2e-07 | 0.44 | 6.9e-06 | 28,1* | Paternal chemotherapy; Cancer of long bones, intestinal tract, lung (secondary): Drugs unknown |

### Paternal defects in DNA repair

For eight of the twelve individuals, the DNMs phased paternally significantly more than expected given the overall ratio of paternal:maternal mutation in the 100kGP cohort (p<0.05/12, Binomial test, Table 1). This implicates the paternal germline as the source of the hypermutation. Two of these fathers carry rare homozygous nonsynonymous variants in known DNA repair genes (Supplemental Table 3). Defects in DNA repair are known to increase the mutation rate in the soma and may have a similar effect in the germline. GEL_1 has the largest number of DNMs of all individuals, a 6.5-fold enrichment, and a significantly increased number of dnIndels. The mutational spectra demonstrates a high enrichment of C>A and T>A mutations (Figure 1b) and we observed a large contribution from Somatic Mutational Signature 8 (Figure 2a). This signature is associated with transcription-coupled nucleotide excision repair (TC-NER) and typically presents with transcriptional strand bias. This agrees with the strong transcriptional strand bias observed in GEL_1 (p = 2.1 ×10^-40^, Poisson test, Supplemental Figure 6). The father has a rare homozygous nonsense variant in the gene *XPC* (Table 1, Supplemental Table 3) which is involved in the early stages of the nucleotide-excision repair (NER) pathway. The paternal variant is annotated as pathogenic for xeroderma pigmentosum in ClinVar and clinical follow-up confirmed that the father had already been diagnosed with this disorder. Patients with xeroderma pigmentosum have a high risk of developing skin cancer due to their impaired ability to repair UV damage and are also known to be at a higher risk of developing other cancers^41,42^. *XPC* deficiency has been associated with a similar mutational spectrum to the one we observe in GEL_1^43^ and xpc deficiency in mice has been shown to increase the germline mutation rate at two STR loci^44^.

GEL_3 has a ~5-fold enrichment of the number of dnSNVs. These dnSNVs exhibit a very distinct mutational spectrum with a ~17-fold increase in T>C mutations but no significant enrichment for any other mutation type (Figure 2b, Supplemental Figure 5d). Extraction of mutational signatures revealed that the majority of mutations mapped onto Somatic Mutational Signature 26 which has been associated with defective mismatch repair. The father has a rare homozygous missense variant in the gene *MPG* (Table 1, Supplemental Table 4). *MPG* encodes N-methylpurine DNA glycosylase (also known as alkyladenine-DNA glycosylase – AAG) which is involved in the recognition of base lesions, including alkylated and deaminated purines, and initiation of the base-excision repair (BER) pathway. The *MPG* variant is rare in gnomAD (allele frequency=9.8 ×10^-5^, no observed homozygotes) and is predicted to be pathogenic by the Combined Annotation Dependent Depletion (CADD) score (CADD score = 27.9) and the amino acid residue is fully conserved across 172 aligned protein sequences from VarSite^45,46^. In the context of the protein, the variant amino-acid forms part of the substrate binding pocket and likely affects substrate specificity (Figure 3a). *MPG* has not yet been described as a cancer susceptibility gene, but studies in yeast and mice have demonstrated variants in this gene, and specifically the substrate binding pocket, can lead to a mutator phenotype^47,48^. We explored the functional impact of the observed A135T variant using *in vitro* assays (Methods, Supplemental Figures 8, 9). The A135T variant caused a two-fold decrease in excision efficiency of the deamination product hypoxanthine (Hx) in both the T and C contexts (Figure 3c, Supplemental Figure 9), with a small increase in excision efficiency of an alkylated adduct 1,N6-ethenoadenine (εA) in both the T and C contexts (Figure 3b, Supplemental Figure 9). The maximal rate of excision is increased by 2-fold for εA which is among the largest increases that have been observed for 15 reported MPG variants (Supplemental Table 4). Another variant, N169S, which also shows an increase in N-glycosidic bond cleavage with the εA substrate has been established as a mutator in yeast^48,49^. These assays confirm that the A135T substitution alters the MPG binding pocket and changes the activity towards different DNA adducts. MPG acts on a wide variety of DNA adducts and further functional characterisation and mechanistic studies are required to link the observed T>C germline mutational signature to the aberrant processing of a specific class of DNA adducts.

**Figure 3.**
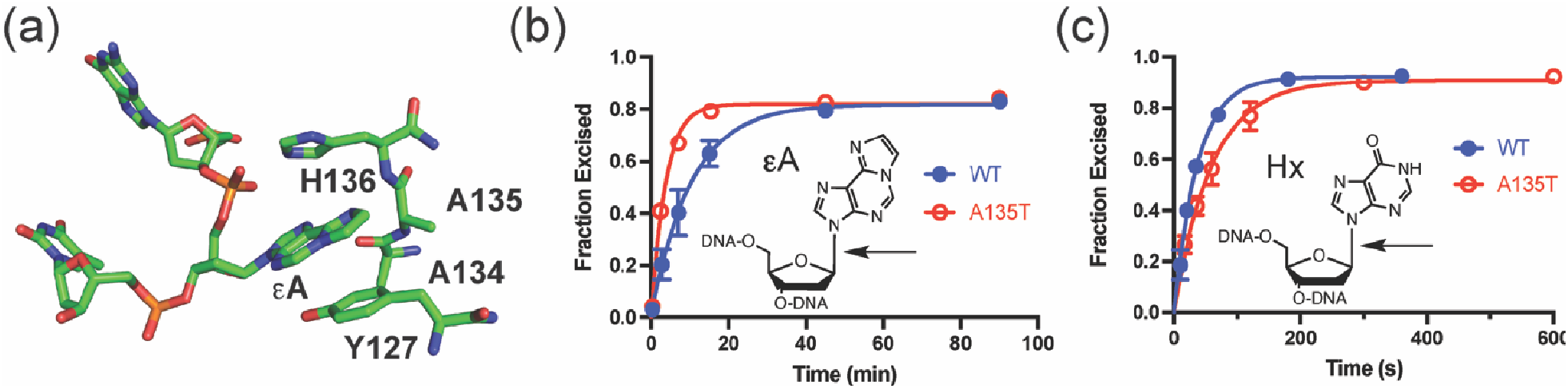
A135T substitution alters the DNA glycosylase activity of MPG. (a) Active site view of MPG bound to εA-DNA from pdb 1EWN. A135 and H136 form the binding pocket for the flipped-out base lesion, which is bracketed by Y127 on the opposing face. (b) Single turnover excision of *εA* from εA·T is 2 fold faster for A135T (red) than for WT (blue) MPG. (c) Single turnover excision of Hx from Hx·T is slower for A135T (red) as compared to WT (blue) MPG. Arrows indicate the N-glycosidic bond that is cleaved by MPG. Each data point is the mean ± SD (N≥3) (see Supplemental Figure 9 for complete kinetic analysis).

GEL_6 has 164 dnSNVs and has a larger contribution of paternally phased mutations than expected (38:3 paternal:maternal, p = 0.022, Binomial test) however the father does not have any nonsynonymous variants in DNA repair genes nor has undergone any chemotherapeutic treatment. This unexplained cause of hypermutation could be due to a paternal variant in a gene currently not associated with DNA repair, a paternal mutator variant that is only present in the germline and not the blood, or a rare gene-by-environment interaction.

### Parental treatment with chemotherapy prior to conception

Three hypermutated individuals (GEL_8, GEL_9 and GEL_11) have a contribution from somatic mutational signature 31 (SBS31) (Figure 2a), which has been associated with treatment with platinum-based drugs such as cisplatin^16^. The phased dnSNVs in GEL_9 and GEL_11 are paternally biased (46 paternal: 2 maternal, p = 0.0014; 28 paternal:1 maternal, p = 0.012; Binomial test, Table 1), and the dnSNVs in GEL_11, which has the largest contribution of SBS31, exhibit a significant transcriptional strand bias, as expected for this signature (p = 6.9 ×10^-6^, Table 1, Supplemental Figure 6). All three fathers have a history of cancer and chemotherapeutic treatment prior to conception as recorded in their available hospital episode records. The father of GEL_11 was diagnosed and received chemotherapeutic treatment for osteosarcoma, lung cancer, and cancer of the intestinal tract within 5 years prior to conception. Cisplatin is a commonly used chemotherapeutic for osteosarcoma and lung cancer. Platinum-based drugs damage DNA by causing covalent adducts. Cisplatin mainly reacts with purine bases, forming intrastrand crosslinks which can be repaired by NER or bypassed by translesion synthesis which may, in turn, induce single base substitutions^50^. The fathers of GEL_8 and GEL_9 both have a history of testicular cancer where cisplatin is the most commonly administered chemotherapeutic.

GEL_2 and DDD_1 have a similar number of dnSNVs, which are significantly paternally biased (Table 1). The mutational spectra of the DNMs in these individuals are very similar and share a novel mutational signature (SBSHYP) that is characterised by an enrichment of C>G and T>G mutations (Figure 2a,b) and does not map on to any previously described signature observed in somatic mutations (as described in COSMIC) or in response to mutagenic exposure^24,51,52^ (Supplemental Figure 10a). The fathers of these individuals do not have putative damaging variants in any DNA repair genes and do not share rare nonsynonymous variants in any other gene. Both fathers received chemotherapeutic treatment prior to conception including nitrogen mustard alkylating agents (Supplemental Table 4), although with different members of this class of chemotherapies, therefore we strongly suspect this class of chemotherapies to be the cause of this novel mutational signature. Experimental studies of a subset of alkylating agents have shown them to have diverse mutational signatures^24,51,52^ (Supplemental Figure 10b).

GEL_5 has 182 dnSNVs and a significant paternal bias in the phased dnSNVs (p = 5.8×10^-4^, Binomial Test, Table 1). The father of GEL_5 has a diagnosis of Systemic Lupus Erythematosus (SLE) and received a course of chemotherapy nine years prior to the conception of the child however the dnSNVs do not map onto any known chemotherapeutic mutational signatures (Figure 1b, Figure 2a). There is a contribution of SBS24 which is associated with aflatoxin exposure in cancer blood samples, however there is no evidence of exposure in the father’s hospital records^22^

We assessed how parental cancer and exposure to chemotherapy might impact germline mutation rate more generally by systematically examining hospital episode statistics across the 100kGP cohort for ICD10 codes related to cancer and chemotherapy that were recorded prior to the conception of the child. We identified 27 fathers (0.9%) who had a history of cancer, 7 of which had testicular cancer (Supplemental Table 6). The offspring of these 27 fathers did not have a significantly increased number of dnSNVs after correcting for parental age (p = 0.73, Wilcox test). This is a small number of fathers so this is not well powered and it is not known definitively how many of these fathers were treated with chemotherapy (6 of the 27 had chemotherapy-related ICD10 codes). Treatment exposure may predate the availability of digitised hospital records and there was limited information on whether conception may have been achieved using sperm stored prior to treatment. While the total number of dnSNVs across all the children is not significantly increased, two of the 27 fathers had hypermutated children which is a significant enrichment compared to those fathers who do not have a recorded history of cancer (2/27 vs 9/2891, p = 0.0043, Fisher exact test). This is likely a conservative p-value as we know that two other hypermutated individuals have fathers who have been treated with chemotherapy however did not fall into this group due to the filtering criteria as we only considered fathers who had at least one ICD 10 code recorded prior to the child’s conception (see Methods). A possible confounder could be that fathers who had cancer prior to conception are older and may be more likely to have been exposed to other germline mutagens however these two groups had the same median paternal age (p = 0.77, Wilcoxon test).

We performed the same analysis across 5,508 mothers in the 100kGP cohort who had hospital episode records entered prior to the conception of their child and identified 27 mothers (0.5%) who had a history of cancer, 9 of whom also had recorded chemotherapy codes. Children whose mothers had a history of cancer had a nominally significant increase of dnSNVs after correcting for parental age and data quality (p = 0.03, Wilcox Test). Mothers who had been diagnosed with cancer were significantly older at the birth of the child compared to those who were not (p = 0.003, Wilcoxon test). Matching on parental age, mothers who had a cancer diagnosis prior to conception had a median increase of 9 dnSNVs. Overall there is not an excess of maternally phased DNMs across these individuals (p = 0.44, Binomial test) however there is one individual with nominal significance (MatCancer_23, 22 paternal:14 maternal, p =0.02, Binomial test, Supplemental Table 5).

We extracted mutational signatures for all these offspring with a maternal or paternal history of cancer that were not hypermutated and found that only one individual had unusual mutational signatures (Supplemental Figure 11).This individual (PatCancer_10) has a contribution of mutational signature SBS31 which is associated with treatment with platinum-based drugs (Supplemental Figure 11). Their father was treated for testicular cancer prior to conception and the child has 94 dnSNVs (p = 0.005, SNV p-value after correcting for parental age) of which 89% phased paternally (p = 0.12, Binomial test, Supplemental Figure 11, Supplemental Table 6).

### Post-zygotic hypermutation

The two hypermutated individuals, GEL_4 and GEL_7, have a ~4 fold and ~2 fold increase in dnSNVs respectively that phase equally between the maternal and paternal chromosomes. The allele balance of the dnSNVs in these individuals was shifted below 0.5 (Supplemental Figure 7). In both individuals, the proportion of DNMs with variant allele fraction (VAF) <0.4 was significantly higher compared to all DNMs across all individuals (GEL_4: p = 3.9 ×10^-59^, GEL_7: p = 8.3×10^-4^, Binomial test). These observations indicate that these mutations most likely occurred post-zygotically and are less likely due to a parental hypermutator. Both individuals have a large contribution of mutations from Somatic Mutational Signature 1 (Figure 2a)^40^. The observations in GEL_4 are likely due to clonal haematopoiesis leading to a large number of somatic mutations in the child’s blood. The mutational signature associated with haematopoietic stem cells is similar to SBS1and we identified a mosaic *de novo* missense mutation in the gene *ETV6.* Mutations in *ETV6* are associated with Leukemia and Thrombocytopenia^53^. GEL_4 has several blood related clinical phenotypes such as abnormality of blood and blood-forming tissues and myelodysplasia. We do not observe similar phenotypes in GEL_7, nor did we identify a possible genetic driver of clonal haematopoiesis, and the child was one year old at recruitment. For this individual we considered the possibility that a maternal mutator variant protein may be impacting the mutation rate in the first few cell divisions. We identified a maternal mosaic missense variant in *TP53* which is annotated as pathogenic in ClinVar for Li-Fraumeni syndrome which is characterised by a predisposition to cancer however it is unknown if this variant is also present in the maternal germline and, if present, whether it would be likely to have a germline mutagenic effect^54^. This variant is not observed in the child.

### Fraction of germline mutation rate variation explained

We investigated the factors influencing the number of dnSNVs per individual in a subset of 7,700 100kGP trios filtered more stringently for data quality (Methods). Using a negative binomial model, accounting for the underlying Poisson variation in germline mutation rate, we estimated that parental age accounts for 69.7% and data quality metrics (eg. read depth, proportion of mapped reads) explain 1.3% of the variance. The variance explained by parental age is smaller than a previous estimate of 95% based on a sample of 78 families^3^. To assess whether this could be due to uncertainty in the previous estimate, we performed repeated sampling of 78 trios from the 100kGP and refit the model and found that the estimates of the variance explained by parental age can vary dramatically with this smaller sample size (median of 79%, 95% interval [52-100%]) and that 7% of our simulations had an estimate as or more extreme than 95%.

We extended this model to account for germline hypermutation by including a variable for the number of excess dnSNVs in the 11 hypermutated individuals in this cohort. We found this explained an additional 7.1% of variance. This leaves 21.9% (19.7%, 23.8%, Bootstrap 95% confidence interval) of variance for numbers of dnSNV per individual unaccounted for. Both mutagenic exposures and genetic variation in DNA repair genes are implicated here as causes of hypermutation, therefore they may also play a more subtle role in the remaining germline mutation rate variation. In addition, polygenic effects and gene by environment interactions may also contribute.

To assess whether rare variants in genes known to be involved in DNA repair pathways impact germline mutation rate more generally, we looked across the whole 100kGP cohort. We curated three sets of rare nonsynonymous variants that have increasing likelihoods of impacting germline mutation rate: (i) variants in all DNA repair genes (N=186), (ii) variants in genes encoding components of the DNA repair pathways most likely to create SNVs (N=66) and (iii) a subset of these variants that had previously been associated with cancer (see Methods). We focused primarily on the effect of heterozygous variants (MAF< 0.001). In the first set of genes we also considered the impact of rare homozygous variants (MAF<0.01) (the counts were too small to assess in the subsequent groups). There was no statistically significant effect in any of these groups of variants after Bonferroni correction (Supplemental Figure 12, Supplemental Table 7). We examined heterozygous protein-truncating variants (PTVs) in the known cancer mutator gene MBD4 which are associated with a three-fold elevated CpG>TpG mutation rate in tumours. We identified and wholegenome sequenced 13 paternal carriers of MBD4 PTVs from the DDD cohort. We found that these individuals did not have a significant increased number of overall DNMs and there was no significant increase in the number of CpG>TpG mutations (p = 0.56, chi-squared test, Supplemental Figure 13). Power modelling suggested there is unlikely to be more than a 22% increase in the CpG mutation rate. This further demonstrates that heterozygous PTVs in known somatic mutator genes may not always have a similar effect in the germline.

To explore the polygenic contribution to germline mutation rate, we estimated the residual variation in the number of dnSNVs in offspring that was explained by germline variants after correcting for parental age, data quality and hypermutation status. We estimated this separately for fathers and mothers in the 100kGP cohort using GREML-LDMS^55^ stratified by minor allele frequency and LD. We found that maternal germline variation (MAF>0.001) did not explain any residual variation (h^2^ = 0.07, p = 0.21, GCTA reported results, Supplemental Table 8). We found that paternal variation may contribute a substantial fraction of residual variation (h^2^ = 0.53 [0.20,0.85], p = 0.09) however this is concentrated exclusively in low frequency variants (0.001 <MAF<0.01, h^2^ = 0.52 [0.01,0.94]) rather than more common variants (MAF> 0.01, h^2^ = 0.008 [0,0.38], Supplemental Table 7). This will need further investigation with larger sample sizes.

## Discussion

Germline hypermutation is an uncommon but important phenomenon. We identified 12 hypermutated individuals from over 20,000 parent offspring sequenced trios in the DDD and 100kGP cohorts with a 2-7 fold increased number of dnSNVs. It is likely that there are additional, currently undetected, germline hypermutated individuals in the DDD cohort. The stringent strategy we adopted to screen this exome-sequenced cohort for potential hypermutated individuals for subsequent confirmation by genome sequencing will have missed some individuals with hypermutation of 2-7 fold.

In two of the 12 hypermutated individuals, the excess mutations appeared to have occurred post-zygotically, however for the majority (n=8) of these hypermutated individuals, the excess dnSNVs phased paternally implicating the father as the source of this hypermutation. For five of these fathers, characteristic mutational signatures and clinical records of cancer treatment prior to conception strongly implicated the mutagenic influence of two different classes of chemotherapeutics: platinum-based drugs (3 families) and mustard-derived alkylating agents (2 families). We also identified likely paternal mutator variants in two hypermutated families. These were rare homozygous missense variants in two known DNA repair genes: *XPC* and *MPG.* Functional and clinical data strongly supported the mutagenic nature of these variants.

It is well established that defects in DNA repair genes can increase somatic mutation rates and elevate cancer risk^56^. Our findings imply that germline mutation rates can be similarly affected. However, defects in DNA repair pathways do not always behave similarly in the soma and the germline. We interrogated PTVs in an established somatic mutator gene, *MBD4,* and found they did not have a detectable effect in the germline^57^. We also examined the impact of parental rare nonsynonymous variants in DNA repair genes on the number of DNMs in offspring and did not find a significant difference. To detect more subtle effects of these variants other analytical approaches will need to be explored. Paternal variants that have previously been associated with a cancer phenotype were nominally significant but having one of these variants only amounted to an estimated average increase of ~2 DNMs in the child. If only a subset of these variants have an impact in the germline this would dilute the power to detect a mutagenic effect and it is likely that both larger sample sizes and additional variant curation will be needed to investigate this further. There may also be genes and pathways that impact mutation in the germline more than the soma; uncovering the genes and associated variants in these genes will be more challenging.

Germline hypermutation accounted for 7% of the variance in germline mutation rate in the 100kGP rare disease cohort. The ascertainment in this cohort for rare disease in the offspring, together with the causal contribution that germline mutation plays in rare diseases, means that germline hypermutated individuals are likely enriched in this cohort relative to the general population. As a consequence, our estimate of the contribution of germline hypermutation to the variance in numbers of dnSNVs per individual is likely inflated. However, the absolute risk of a germline hypermutator having a child with a genetic disease is still low. The population average risk for having a child with a severe developmental disorder caused by a *de novo* mutation has been estimated to be 1 in 300 births^11^ and so a 4-fold increase in DNMs in a child would only elevate this absolute risk to just over 1%. Therefore, we anticipate that most germline hypermutated individuals will not have a rare genetic disease, and germline hypermutation will also be observed in healthy population cohorts.

The two genetic causes of germline hypermutation that we identified were both recessive in action. Similarly, most DNA repair disorders act recessively in their cellular mutagenic effects. This implies that genetic causes of germline hypermutation are likely to arise at substantially higher frequencies in populations with high rates of parental consanguinity. In such populations, the overall incidence of germline hypermutation may be higher and the proportion of the variance in the number of dnSNVs per offspring accounted for genetic effects will be higher. We anticipate that studies focused on these populations are likely to identify additional mutations that affect germline mutation rate.

We found that, among 7,700 100kGP families, parental age only explained ~70% of the variance in numbers of dnSNVs per offspring, which is substantially smaller than a previous estimate of 95% based on a sample of 78 families^3^. Repeated sampling of 78 trios from the 100kGP showed that estimates of the variance explained by parental age can vary dramatically stochastically and we regard our estimate based on two orders of magnitude more trios to be more reliable, although other differences between the studies such as measurement error and criteria for ascertainment of families might be having a subtle influence. The residual ~20% of variation in numbers of germline dnSNVs per individual remains unexplained by parental age, data quality and hypermutation. We found that rare variants in known DNA repair genes are unlikely to account for a large proportion of this unexplained variance. Heritability analyses suggested that polygenic contributions from common variants (MAF>1%) are unlikely to make a substantive contribution to this variance; however, we observed some evidence that the polygenic contribution of intermediate frequency paternal variants (0.001<MAF<0.01) could be more substantial although larger sample sizes are required to confirm this observation. A limitation to these heritability analyses is that we use DNMs in offspring as a proxy for individual germline mutation rates. Measuring germline mutation rates more directly by, for example, sequencing hundreds of single gametes per individual, should facilitate better powered association studies and heritability analyses.

Environmental exposures are also likely to contribute to germline mutation rate variation. We have observed evidence that certain chemotherapeutics can affect germline mutation rate and targeted studies on the germline mutagenic effects of different chemotherapeutics (e.g. in cancer survivor cohorts) will be crucial in understanding this further. We anticipate that these studies will identify considerable heterogeneity in the germline mutagenic effects of different chemotherapeutics, in part due to differences in the pemeability of the blood-testis barrier to different agents^58^, as well as variation in the vulnerability to chemotherapeutic germline mutagenesis by sex and age. As so few individuals are treated for cancer prior to reproduction, chemotherapeutic exposures will not explain a large proportion of the remaining variation in germline mutation rates however chemotherapeutic mutagenesis has important implications for cancer patients who plan to have children, especially in whether they decide to store unexposed gametes for future use of assisted reproductive technologies.

Unexplained hypermutation and additional variance in germline mutation rate may be explained by other environmental exposures. A limitation of this study was the lack of data on non-therapeutic environmental exposures. However, and somewhat reassuringly, the relatively tight distribution of DNMs per person in 100kGP suggests that there are unlikely to be common environmental mutagen exposures in the UK (e.g. cigarette smoking) that causes a substantive (e.g, >1.5 times) fold increase in mutation rates and concomitant disease risk. The germline generally appears to be well protected from large increases in mutation rate. However, including a broader spectrum of environmental exposures in future studies would help to identify more subtle effects and may reveal gene-by-environment interactions.

## Acknowledgments

We thank the families and their clinicians for their participation and engagement, and our colleagues who assisted in the generation and processing of data. We would like to thank Mike Stratton, Peter Campbell, Emily Mitchell, Eleanor Dunstone, Hilary Martin, Kartik Chundru, Molly Przeworski and Jan Korbel for helpful discussions and advice. This research was made possible through access to the data and findings generated by the 100,000 Genomes Project. The 100,000 Genomes Project is managed by Genomics England Limited (a wholly owned company of the Department of Health and Social Care). The 100,000 Genomes Project is funded by the National Institute for Health Research and NHS England. The Wellcome Trust, Cancer Research UK and the Medical Research Council have also funded research infrastructure. The 100,000 Genomes Project uses data provided by patients and collected by the National Health Service as part of their care and support. The DDD study presents independent research commissioned by the Health Innovation Challenge Fund (grant number HICF-1009-003). The full acknowledgements can be found at www.ddduk.org/access.html. This research was funded in part by the Wellcome Trust grant [206194]. For the purpose of open access, the author has applied a CC BY public copyright licence to any Author Accepted Manuscript version arising from this submission. This work was supported by Health Data Research UK, which is funded by the UK Medical Research Council, Engineering and Physical Sciences Research Council, Economic and Social Research Council, Department of Health and Social Care (England), Chief Scientist Office of the Scottish Government Health and Social Care Directorates, Health and Social Care Research and Development Division (Welsh Government), Public Health Agency (Northern Ireland), British Heart Foundation and Wellcome.

## Methods

### DNM filtering in 100,000 Genomes Project

We analysed DNMs called in 13,949 parent offspring trios from 12,609 families from the rare disease programme of the 100,000 Genomes Project. The rare disease cohort includes individuals with a wide array of diseases including neurodevelopmental disorders, cardiovascular disorders, renal and urinary tract disorders, ophthalmological disorders, tumour syndromes, ciliopathies and others. These are described in more detail in previous publications^59,60^. The cohort was whole genome sequenced at ~35X coverage and variant calling for these families was performed via the Genomics England rare disease analysis pipeline. The details of sequencing and variant calling have been previously described^60^. DNMs were called by the Genomics England Bioinformatics team using the Platypus variant caller^61^. These were selected to optimise various properties including the number of DNMs per person being approximately what we would expect, the distribution of the VAF of the DNMs to be centered around 0.5 and the true positive rate of DNMs to be sufficiently high as calculated from examining IGV plots. The filters applied were as follows:

- Genotype is heterozygous in child (1/0) and homozygous in both parents (0/0)
- Child RD >20, Mother RD>20, Father RD>20
- Remove variants with >1 alternative read in either parent
- VAF>0.3 and VAF<0.7 for child
- Remove SNVs within 20 bp of each other. While this is likely removing true MNVs, the error mode was very high for clustered mutations.
- Removed DNMs if child RD >98^13^
- Removed DNMs that fell within known segmental duplication regions as defined by
- UCSC (http://humanparalogy.gs.washington.edu/build37/data/GRCh37GenomicSuperDup.tab)
- Removed DNMs that fell in highly repetitive regions (http://humanparalogy.gs.washington.edu/build37/data/GRCh37simpleRepeat.txt)
- For DNM calls that fell on the X chromosome these slightly modified filters were used:

∘ For DNMs that fell in PAR regions, the filters were unchanged from the autosomal calls apart from allowing for both heterozygous (1/0) and hemizygous (1) calls in males
∘ For DNMs that fell in non-PAR regions the following filters were used:

▪ For males: RD>20 in child, RD>20 in mother, no RD filter on father
▪ For males: the genotype must be hemizygous (1) in child and homozygous in mother (0/0)
▪ For females: RD>20 in child, RD>20 in mother, RD>10 in father

### DNM filtering and identifying hypermutated individuals in DDD

To identify hypermutated individuals in the DDD study we started with exome sequencing data from the DDD study of families with a child with a severe, undiagnosed developmental disorder. The recruitment of these families has been described previously^62^: families were recruited at 24 clinical genetics centers within the UK National Health Service and the Republic of Ireland. Families gave informed consent to participate, and the study was approved by the UK Research Ethics Committee (10/H0305/83, granted by the Cambridge South Research Ethics Committee, and GEN/284/12, granted by the Republic of Ireland Research Ethics Committee). Sequence alignment and variant calling of SNV and insertions/deletions were conducted as previously described. De novo mutations were called using DeNovoGear and filtered as previously^63,11^. The analysis in this paper was conducted on a subset (7,930 parent offspring trios) of the full current cohort which was not available at the start of this research.

In the DDD study, we identified 9 individuals out of 7,930 parent-offspring trios with an increased number of exome DNMs after accounting for parental age (7-17 exome DNMs compared to an expected number of ~2). These were subsequently submitted along with their parents for PCR-free whole-genome sequencing at >30x mean coverage using Illumina 150bp paired end reads and in house WSI sequencing pipelines. Reads were mapped with bwa (v0.7.15)^64^. DNMs were called from these trios using DeNovoGear^63^ and were filtered as follows:

- Read depth (RD) of child > 10, mother RD > 10, father RD > 10
- Alternative allele read depth in child >2
- Filtered on strand bias across parents and child (p-value >0.001, Fisher’s exact test)
- Removed DNMs that fell within known segmental duplication regions as defined by
- UCSC (http://humanparalogy.gs.washington.edu/build37/data/GRCh37GenomicSuperDup.tab)
- Removed DNMs that fell in highly repetitive regions (http://humanparalogy.gs.washington.edu/build37/data/GRCh37simpleRepeat.txt)
- Allele frequency in gnomAD < 0.01
- VAF <0.1 for both parents
- Removed mutations if both parents have >1 read supporting the alternative allele
- Test to see if VAF in child is significantly greater than the error rate at that site as defined by error sites estimated using Shearwater^65^.
- Posterior probability from DeNovoGear > 0.00781^63,11^
- Removed DNMs if child RD >200.

After applying these filters, this resulted in 1,367 DNMs. All of these DNMs were inspected in the Integrative Genome Viewer^66^ and removed if they appeared to be false positives. This resulted in a final set of 916 DNMs across the 9 trios. One of the 9 had 277 dnSNVs genome wide while the remaining had expected numbers (median number of 81 dnSNVs).

### Parental phasing of *de novo* mutations

To phase the DNMs in both 100kGP and DDD we used a custom script which used the following read-based approach to phase a DNM. This first searches for heterozygous variants within 500 bp of the DNM that was able to be phased to a parent (so not heterozygous in both parents and offspring). We then examined the reads or read pairs which included both the variant and the DNM and counted how many times we observed the DNM on the same haplotype of each parent. If the DNM appears exclusively on the same haplotype as a single parent then that was determined to originate from that parent. We discarded DNMs that had conflicting evidence from both parents. This code is available on GitHub (https://github.com/queenjobo/PhaseMyDeNovo).

### Analysis of effect of parental age on germline mutation rate

To assess the effect of parental age on germline mutation rate we ran the following regressions on autosomal DNMs. On all (unphased) DNMs we ran two separate regressions for SNVs and indels. We chose a negative Binomial generalized linear model here as the Poisson was found to be overdispersed. We fitted the following model using a negative Binomial GLM with an identity link where *Y* is the number of DNMs for an individual:

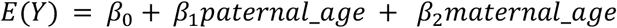

For the phased DNMs we fit the following two models using a negative Binomial GLM with an identity link where *Y_maternal_* is the number of maternally derived DNMs and *Y_paternal_* is the number of paternally derived DNMs:

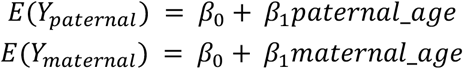

### Identifying hypermutated individuals in 100kGP

To identify hypermutated individuals in the 100kGP cohort we first wanted to regress out the effect of parental age as described in the parental age analysis. We then looked at the distribution of the studentized residuals and then, assuming these followed a *t* distribution with *N-3* degrees of freedom, calculated a t-test p-value for each individual. We took the same approach for the number of indels, except in this case *Y* would be the number of de novo indels.

We identified 21 individuals out of 12,471 parent-offspring trios with significantly increased number of dnSNVs genome wide (p < 0.05/12471). We performed multiple quality control analyses which included examining the mutations in the Integrative Genomics Browser for these individuals to examine DNM calling accuracy, looking at the relative position of the DNMs across the genome and examining the mutational spectra of the DNMs to identify any well known sequencing error mutation types. We identified 12 that were not truly hypermutated. The majority of false positives (10) were due to a parental somatic deletion in blood increasing the number of apparent DNMs (Supplemental Figure 14). These individuals had some of the highest number of DNMs called (up to 1379 DNMs per individual). For each of these 10 individuals, the DNM calls all clustered to a specific region in a single chromosome. In this same corresponding region in the parent, we observed a loss of heterozygosity when calculating the heterozygous/homozygous ratio. In addition, many of these calls appeared to be low level mosaic in that same parent. This type of event has previously been shown to create artifacts in CNV calls and is referred to as a ‘Loss of Transmitted Allele’ event^67^. The remaining 2 false positives were due to bad data quality in either the offspring or one of the parents leading to poor DNM calls. The large number of DNMs in these false positive individuals also led to significant underdispersion in the model so after removing these 12 individuals we reran the regression model and subsequently identified 11 individuals which appeared truly hypermutated (p< 0.05/12,459).

### Extraction of mutational signatures

Mutational signatures were extracted from maternally and paternally phased autosomal DNMs, 24 controls (randomly selected), 25 individuals (father with a cancer diagnosis prior to conception), 27 individuals (mother with a cancer diagnosis prior to conception) and 12 hypermutated individuals that we identified. All DNMs were lifted over to GRCh37 prior to signature extraction (100kGP samples are a mix of GRCh37 and GRCh38) and through the liftover process a small number of 100kGP DNMs were lost (0.09% overall, 2 DNMs lost across all hypermutated individuals). The mutation counts for all the samples can be found in Supplemental Table 1. This was done using SigProfiler (v1.0.17) and these signatures are extracted and subsequently mapped on to COSMIC mutational signatures (COSMIC v91, Mutational Signature v3.1)^19,40^. Sigprofiler defaults to selecting a solution with higher specificity than sensitivity. A solution with 4 de-novo signatures was chosen as optimal by SigProfiler for the 12 hypermutator samples. Another stable solution with five de-novo signatures was also manually deconvoluted, which has been considered as the final solution. The mutation probability for mutational signature SBSHYP can be found in Supplemental Table 2.

### Signature comparison to external exposures

We compared the extracted signatures from these hypermutated individuals to a compilation of previously identified signatures caused by enviromental mutagens from the literature. The environmental signatures were compiled from Kucab et al (Cell 2019)^24^, Pich et al (Nautre Genetics 2019)^51^ and Volkova et al (Nature Communications 2020)^52^. Comparison was calculated as the cosine similarity between the different signatures.

### Defining set of genes involved in DNA repair

We compiled a list of DNA repair genes which were taken from an updated version of the table in Lange et al, Nature Reviews Cancer 2011 (https://www.mdanderson.org/documents/Labs/Wood-Laboratory/human-dna-repair-genes.html)^68^. These can be found in Supplemental Table 3. These are annotated with the pathways they are involved with (eg. nucleotide-excision repair, mismatch repair). A ‘rare’ variant is defined as those with an allele frequency of <0.001 for heterozygous variants and those with an allele frequency of <0.01 for homozygous variants in both 1000 Genomes as well as across the 100kGP cohort.

### Kinetic characterization of MPG

The A135T variant of MPG was generated by site-directed mutagenesis and confirmed by sequencing both strands. The catalytic domain of WT and A135T MPG were expressed in BL21(DE3) Rosetta2 *E. coli* and purified as described for the full-length protein^69^. Protein concentration was determined by absorbance at 280 nm. Active concentration was determined by electrophoretic mobility shift assay with 5’FAM-labeled pyrolidine-DNA (Supplemental Figure 8)^48^. Glycosylase assays were performed with 50 mM NaMOPS, pH 7.3, 172 mM potassium acetate, 1 mM DTT, 1 mM EDTA, 0.1 mg/mL BSA at 37°C. For single turnover glycosylase activity, a 5’-FAM-labeled duplex was annealed by heating to 95°C and slowly cooling to 4°C (see Supplemental Figure 9). DNA substrate concentration was varied between 10 and 50 nM and MPG concentration was maintained in at least 2-fold excess over DNA from 25 to 10,000 nM. Timepoints were quenched in 0.2 M NaOH, heated to 70°C for 12.5 min, then mixed with formamide/EDTA loading buffer and analyzed by 15% denaturing polyacrylamide gel electrophoresis. Fluorescence was quantified with a Typhoon 5 imager and ImageQuant software (GE). The fraction of product was fit by a single exponential equation to determine the observed single turnover rate constant (*k_obs_*). For Hx excision, the concentration dependence was fit by the equation *k_obs_* = *k_max_* [E]/(K_1/2_+[E]), in which the K_1/2_ is the concentration at which half the maximal rate constant (*k_max_*) was obtained and [E] is the concentration of enzyme. It was not possible to measure the K1/2 for *εA* excision using a fluorescence-based assay due to extremely tight binding^70^. Multiple turnover glycosylase assays were performed with 5 nM MPG and 10—40-fold excess of substrate (Supplemental Figure 9).

### Estimating the fraction of variance explained

To estimate the fraction of germline mutation variance explained by several factors, we fit the following negative Binomial GLMs with an identity link. Data quality is likely to correlate with the number of DNMs detected so to reduce this variation we used a subset of the 100kGP dataset which had been filtered on some base quality control (QC) metrics by the Bioinformatics team at GEL:

- cross-contamination < 5%
- mapping rate > 75%
- mean sample coverage > 20
- insert size <250

We then included the following variables to try and capture as much of the residual measurement error which may also be impacting DNM calling. In brackets are the corresponding variable names used in the models below:

- Mean coverage for the child, mother and father *(child_mean_RD, mother_mean_RD, father_mean_RD)*
- Proportion of aligned reads for the child, mother and father *(child_prop_aligned, mother_prop_aligned, father_prop_aligned)*
- Number of SNVs called for child, mother and father (child_snvs, mother_snvs, father_snvs)
- Median VAF of DNMs called in child *(median_VAF)*
- Median ‘Bayes Factor’ as outputted by Platypus for DNMs called in the child. This is a metric of DNM quality *(median_BF).*

The first model only included parental age:

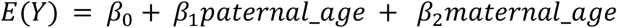

The second model also included data quality variables as described above:

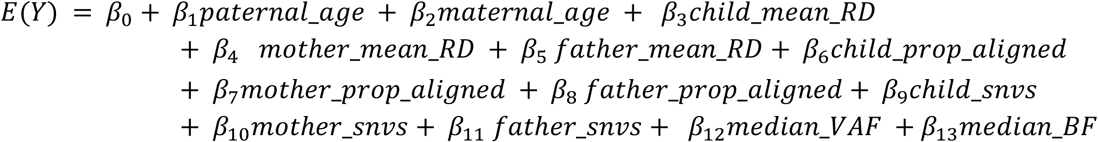

The third model included a variable for excess mutations in the 11 confirmed hypermutated individuals (hm_excess) in the 100kGP dataset. This variable was the total number of mutations subtracted by the median number of DNMs in the cohort (65), Y_hypermutated_ – median(Y) for these 11 individuals and 0 for all other individuals.

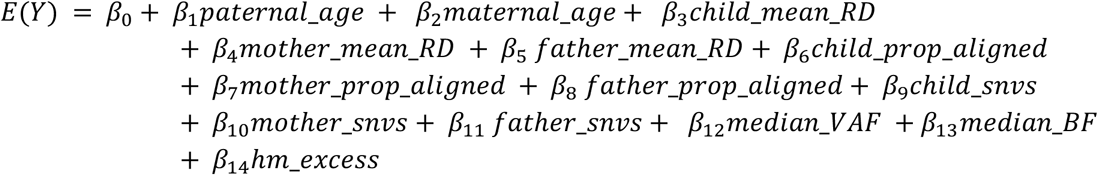

The fraction of variance (*F*) explained after accounting for Poisson variance in the mutation rate was calculated in a similar way to Kong et al using the following formula^3^.

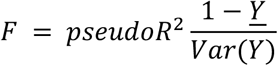

McFadden’s pseudo R^2^ was used here as a Negative binomial GLM was fitted. We repeated these analyses fitting an ordinary least squares regression, as was done in Kong et al^3^, using the R^2^ and got comparable results. To calculate a 95% confidence interval we used a bootstrapping approach. We sampled with replacement 1,000 times and extracted the 2.5% and 97.5% percentiles.

### Analysis of contribution of rare variants in DNA repair genes

We fit 8 separate regressions to assess the contribution of rare variants in DNA repair genes (compiled as described previously). These were across three different sets of genes: variants in all DNA repair genes, variants in a subset of DNA repair genes known to be associated with BER, MMR, NER or a DNA polymerase and variants within this subset that have also been associated with a cancer phenotype. For this we downloaded all ClinVar entries as of October 2019 and searched for germline ‘pathogenic’ or ‘likely pathogenic’ variants annotated with cancer^54^. We tested both all nonsynonymous variants and just protein truncating variants (PTVs) for each set. To assess the contribution of each of these sets we created two binary variables per set indicating a presence or absence of a maternal or paternal variant for each individual and then ran a negative binomial regression for each subset including these as independent variables along with hypermutation status, parental age and QC metrics as described in the previous section.

### Simulations to explore effect estimates of fraction of variance explained by paternal age from downsampling

To explore how the estimates of the fraction of variance of the number of DNMs is explained by paternal age varies with downsampling we first simulated a random sample as follows 10,000 times:

- Randomly sample 78 trios (the number of trios in Kong et al.^3^.)
- Fit OLS of *E*(*Y*) = *β*_0_ + *β*_1_*paternal_age*
- Estimated fraction of variance (*F*) as described in Kong et al^3^.

We found that the median fraction explained was 0.77, sd of 0.13 and with 95% of simulations fallings between 0.51 and 1.00.

### Identifying parents with cancer diagnosis prior to birth of offspring

To identify parents who had received a cancer diagnosis prior to the conception of their child we examined the admitted patient care hospital episode statistics of these parents. There were no hospital episode statistics available prior to 1997 and many individuals did not have any records until after the birth of the child. To ensure comparisons were not biased by this we first subsetted to parents who had at least one episode statistic recorded at least two years prior to the child’s year of birth. Two years prior to the child’s birth was our best approximation for prior to conception without the exact child date of birth. This resulted in 2,891 fathers and 5,508 mothers. From this set we then extracted all entries with ICD10 codes with a “C” prefix which corresponds to malignant neoplasms and “Z85” which corresponds to a personal history of malignant neoplasm. We defined a parent as having a cancer diagnosis prior to conception if they had any of these codes recorded >=2 years prior to the child’s year of birth. We also extracted all entries with ICD10 code “Z511” which codes for an ‘encounter for antineoplastic chemotherapy and immunotherapy’.

Two fathers of hypermutated individuals who we suspect had chemotherapy prior to conception did not meet these criteria as the father of GEL_5 received chemotherapy for treatment for SLE and not cancer and for the father of GEL_8 the hospital record ‘personal history of malignant neoplasm’ were entered after the conception of the child (Supplemental Table 4).

To compare the number of dnSNVs between the group of individuals with parents with and without cancer diagnoses we used a Wilcoxon test on the residuals from the negative binomial regression on dnSNVs correcting for parental age, hypermutation status and data quality. To look at the effect of maternal cancer on dnSNVs we matched these individuals on maternal and paternal age with sampling replacement with 20 controls for each of the 27 individuals. We found a significant increase in DNMs (74 compared to 65 median dnSNVs, p = 0.001, Wilcoxon Test).

### SNP heritability analysis

For this analysis we started with the same subset of the 100kGP dataset that had been filtered as described in the analysis on the impact of rare variants in DNA repair genes across the cohort (see above). To ensure variant quality we subsetted to variants that have been observed in genomes from gnomAD (v3)^71^. These were then filtered by ancestry to parent-offspring trios where both the parents and child mapped on to the 1000 Genomes GBR subpopulations. The first 10 principal components were subsequently included in the heritability analyses. To remove cryptic relatedness we removed individuals with estimated relatedness >0.025 (using GCTA grm-cutoff 0.025). This resulted in a set of 6,352 fathers and 6,329 mothers. The phenotype in this analysis was defined as the residual from the negative binomial regression of the number of DNMs after accounting for parental age, hypermutation status and several data quality variables as described when estimating the fraction of DNM count variation explained (see Methods above). To estimate heritability we ran GCTA’s GREML-LDMS on two LD stratifications and three MAF bins (0.001-0.01,0.01-0.05,0.05-1).^55^ For mothers this was run with the --reml-no-constrain option because otherwise it would not converge. (Supplemental Table 8)

