## Supplemental Figures/Tables for "Genetic and chemotherapeutic causes of germline hypermutation"

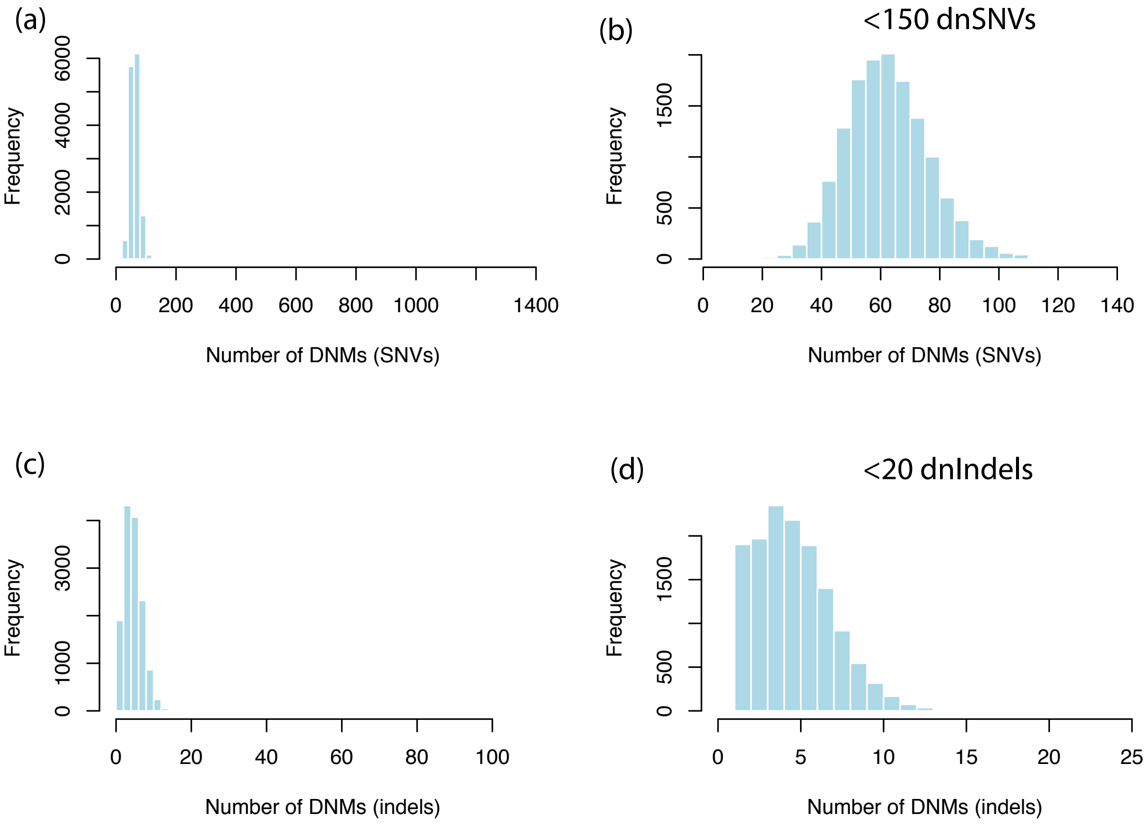


Supplemental Figure 1: Distribution of number of *de novo* SNVs for all individuals (a) and those with <150 DNMs (b). Distribution of number of *de novo* InDels per person for all individuals (c) and those with <20 indels (d).

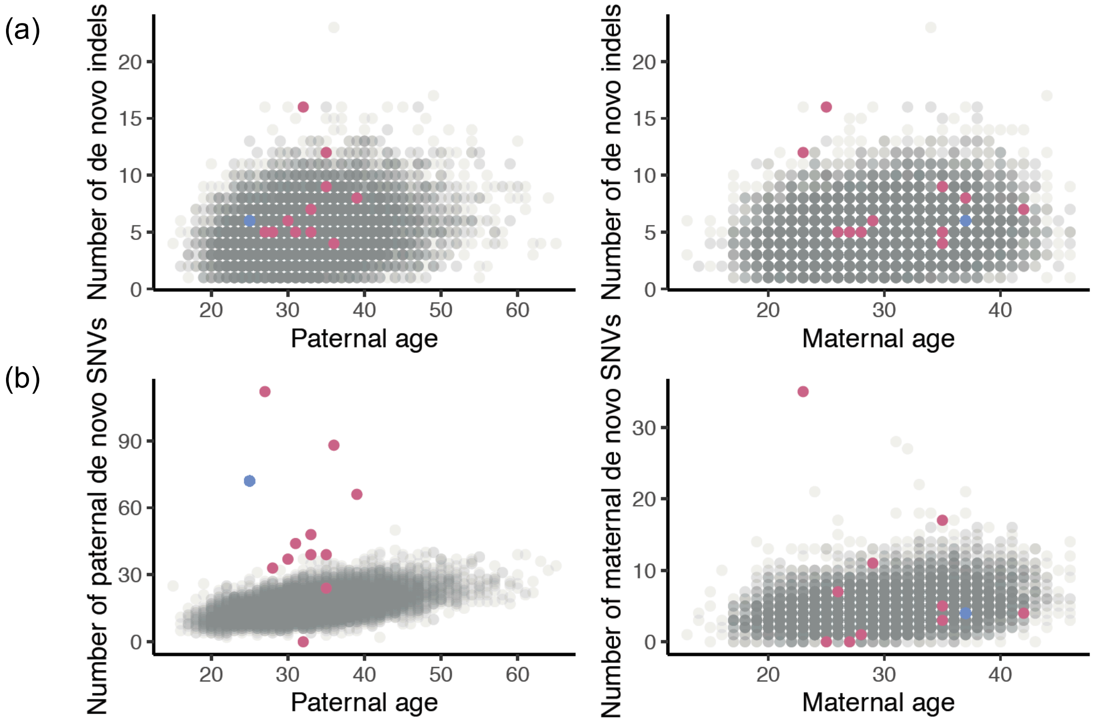


Supplemental Figure 2: Paternal and maternal age against the number of (a) dnInDels. (b) Paternal age against number of paternally phased dnSNVs and maternal age against number of maternally phased dnSNVs. Hypermutated individuals are highlighted in pink (11 individuals in 100kGP) and blue (DDD individuals).

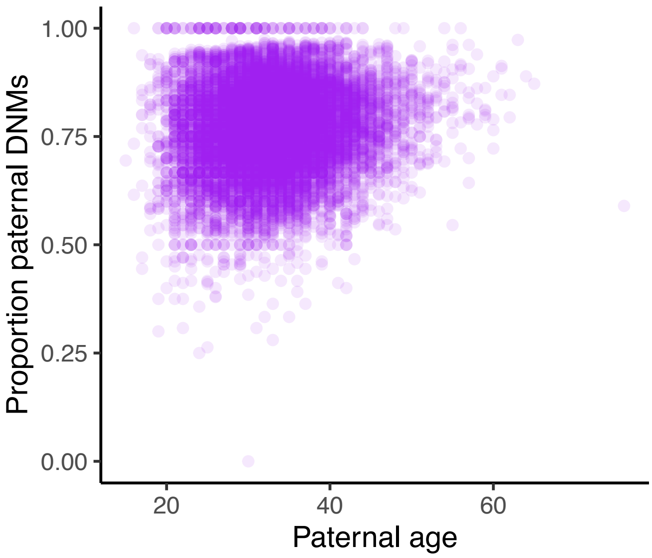


Supplemental Figure 3: Proportion of paternally phased DNMs against paternal age. X-axis refers to paternal age at child's birth. Y-axis is the proportion of phased DNMs that phased paternally.

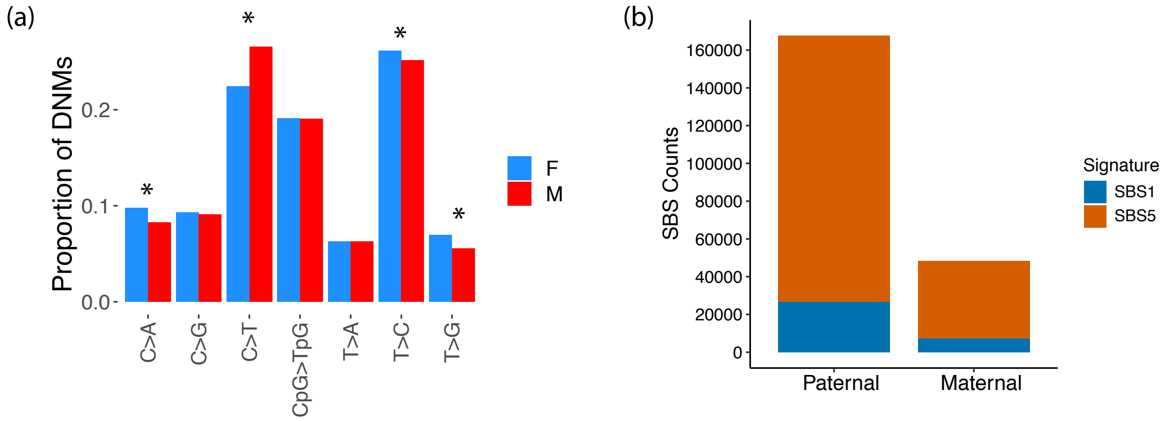


Supplemental Figure 4: Mutational spectra and signatures for maternal vs paternal DNMs across 100kGP cohort (a) Mutational spectra for maternal vs paternal DNMs across 100kGP cohort. Significant differences (p<0.05/7) are marked with *. (b) Mutational signature decomposition for DNMs in maternally and paternally derived DNMs. Signatures extracted with SigProfiler. Colors correspond to COSMIC signatures.

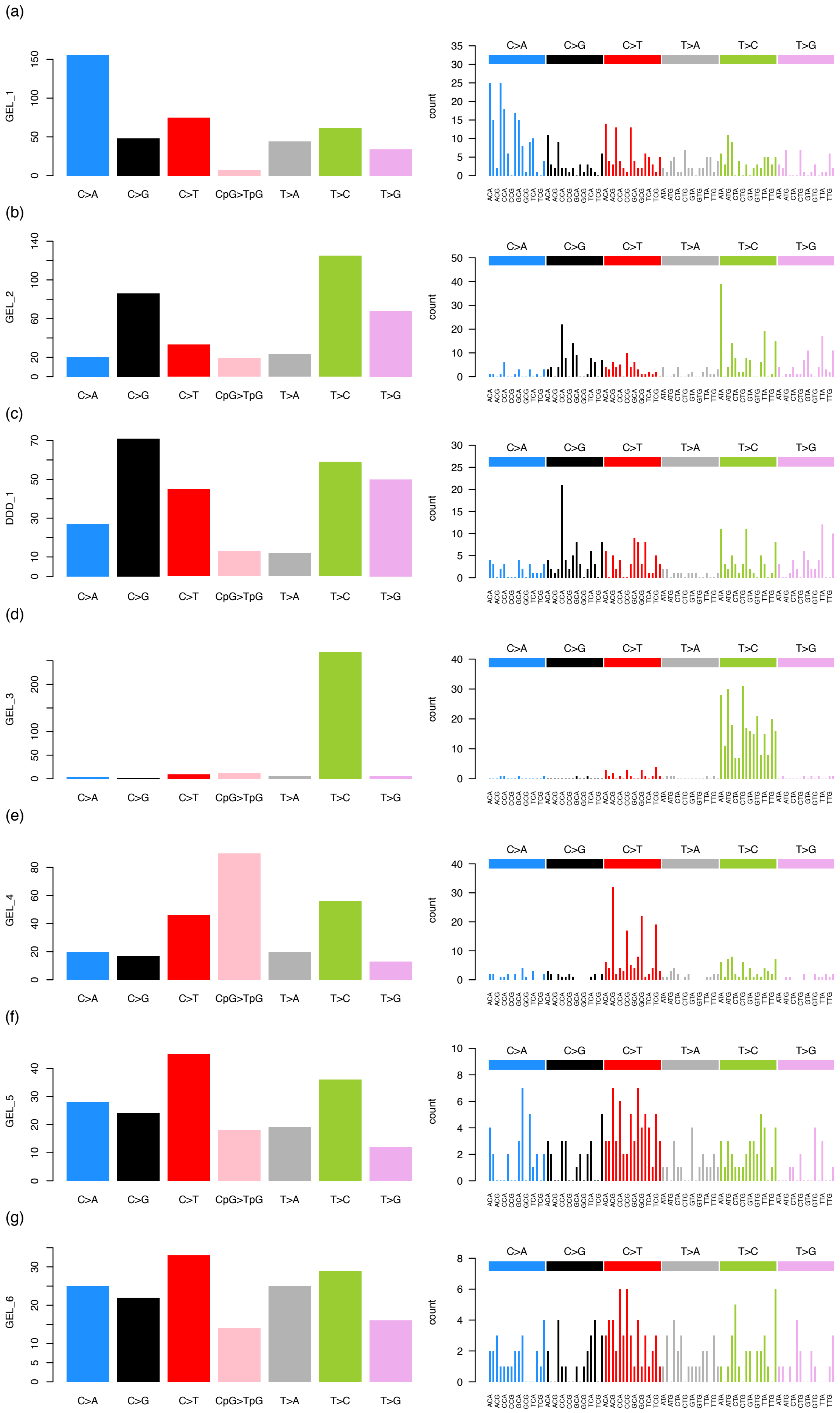

(a)
GEL_1
C>A
C>G
C>T
CpG>TpG
T>A
T>C
T>G
count
(b)
GEL_2
(c)
DDD_1
(d)
GEL_3
(e)
GEL_4
(f)
GEL_5
(g)
GEL_6

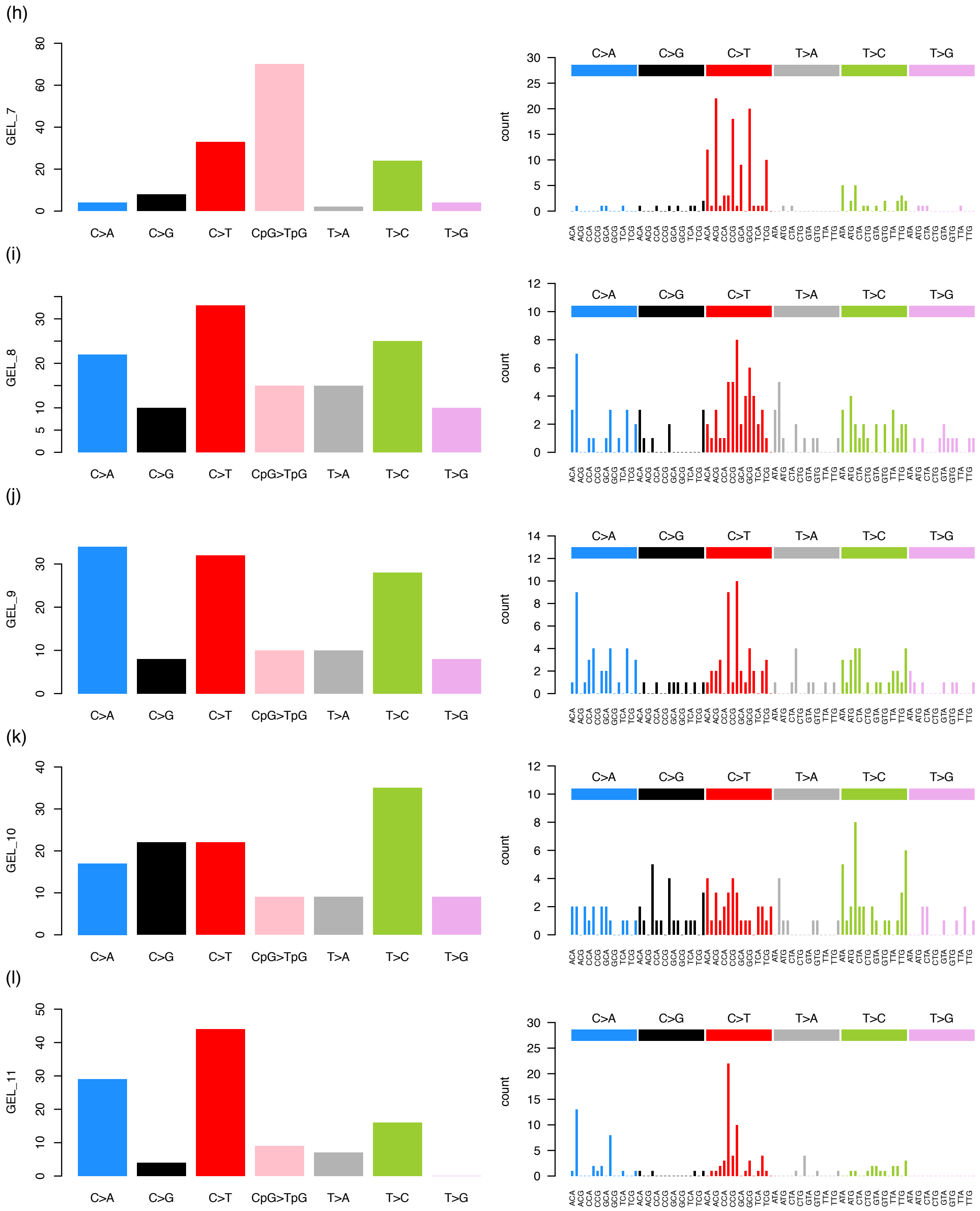


Supplemental Figure 5: Mutational spectra for the DNMs of the 12 hypermutated individuals. Each row is a hypermutated individual showing the mutational spectra according to count of mutations per each single base change (with CpG>TpG mutations separated from other C>T mutations) and the second plot is the mutation count for all 96 mutations in their trinucleotide context. The x-axis here demonstrate the reference trinucleotide sequence with the mutated base highlighted. The color and label on bar above indicates the mutation type.

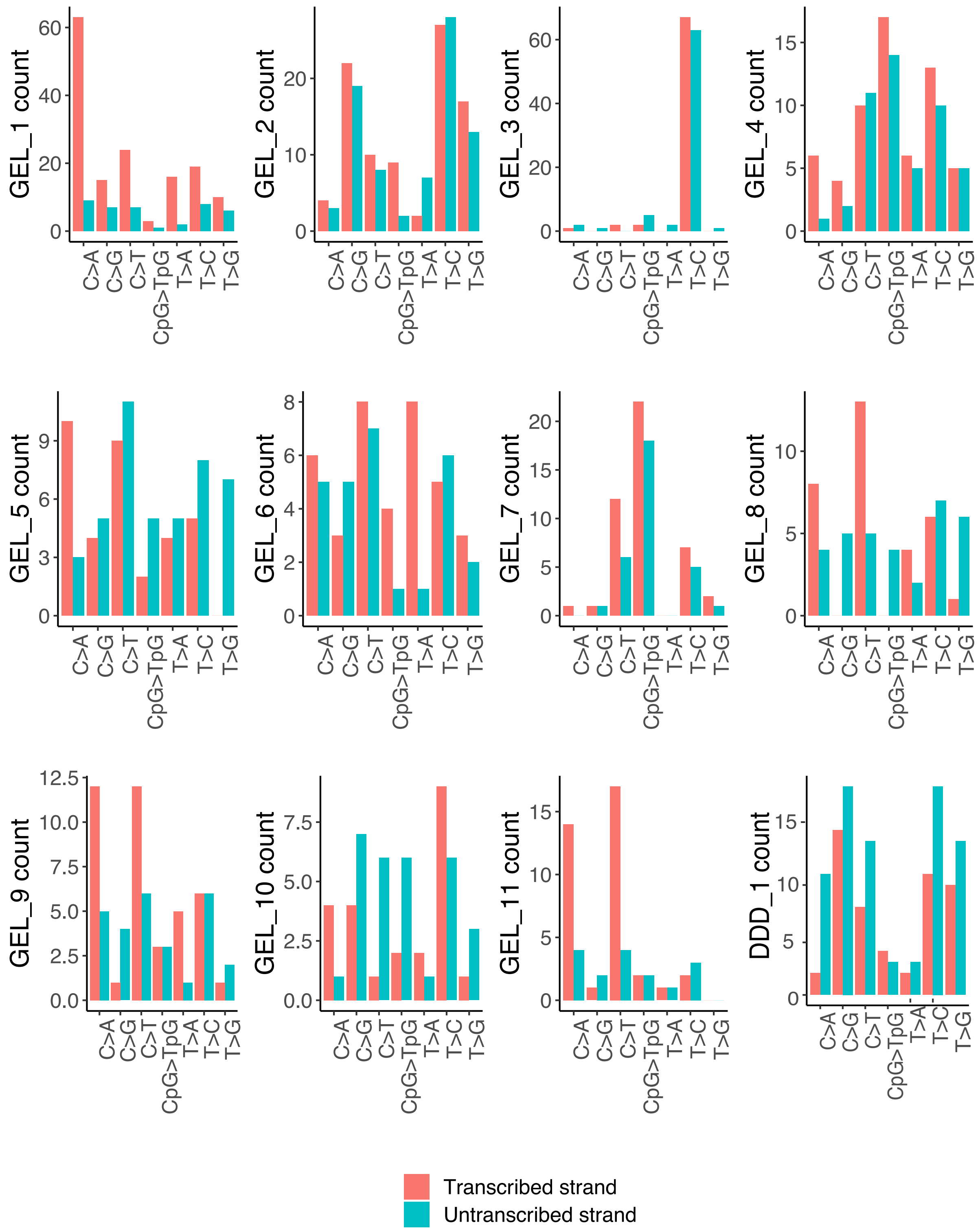


Supplemental Figure 6: Transcriptional strand bias for DNMs in the 12 hypermutated individuals. This demonstrate the count of each mutation type on the transcribed and untranscribed strand for each individual.

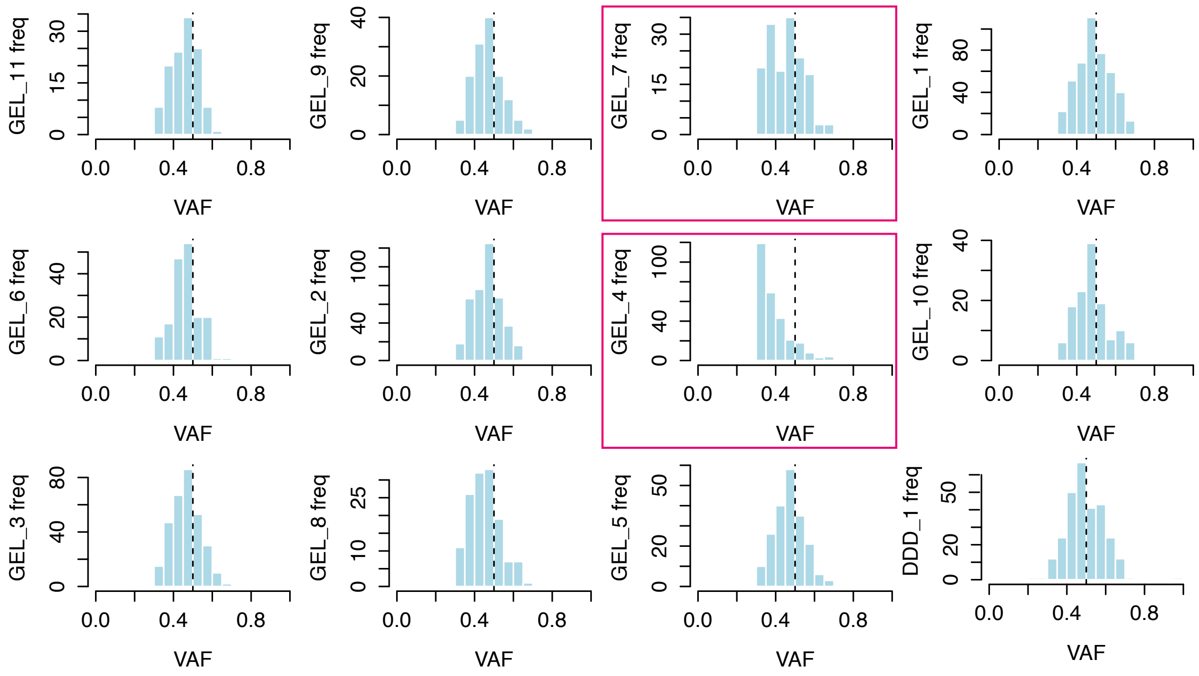


Supplemental Figure 7: Distribution of variant allele fraction (VAF) for DNMs in hypermutated individuals. The vertical line indicates 0.5 VAF. The two plots highlighted in pink are those where the DNMs appear post-zygotic.

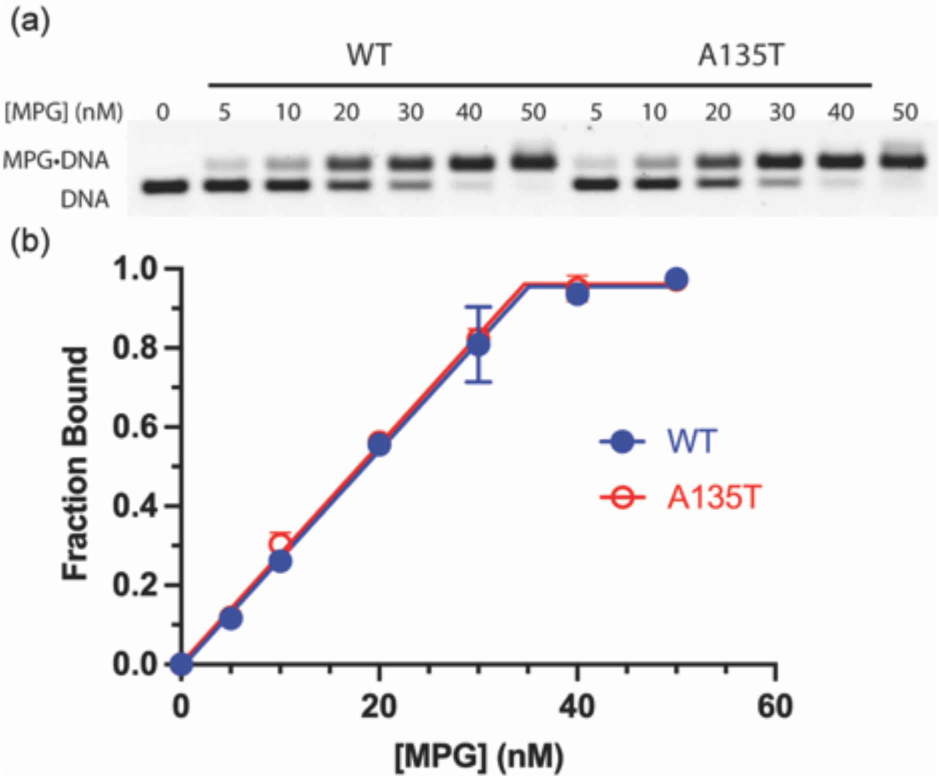


Supplemental Figure 8. Determination of active concentration of MPG. (a) Representative native gel electrophoresis with 20 nM pyrolidine-DNA (Y•T) and varying concentration of WT or A135T MPG (25 mM NaHEPES pH 7.5, 100 mM NaCl, 5% v/v glycerol, 1 mM EDTA, 1 mM DTT). Agarose gels (2% w/v) were run in 0.5X TBE buffer at 10V/cm at 4 oC. (b) Independent dilutions were fit to a binding titration to yield an active fraction of 0.57 for both WT and A135T (N=3). This demonstrates that equal concentrations of WT and A135T were tested in the glycosylase assays. The concentrations listed are not corrected by this factor.

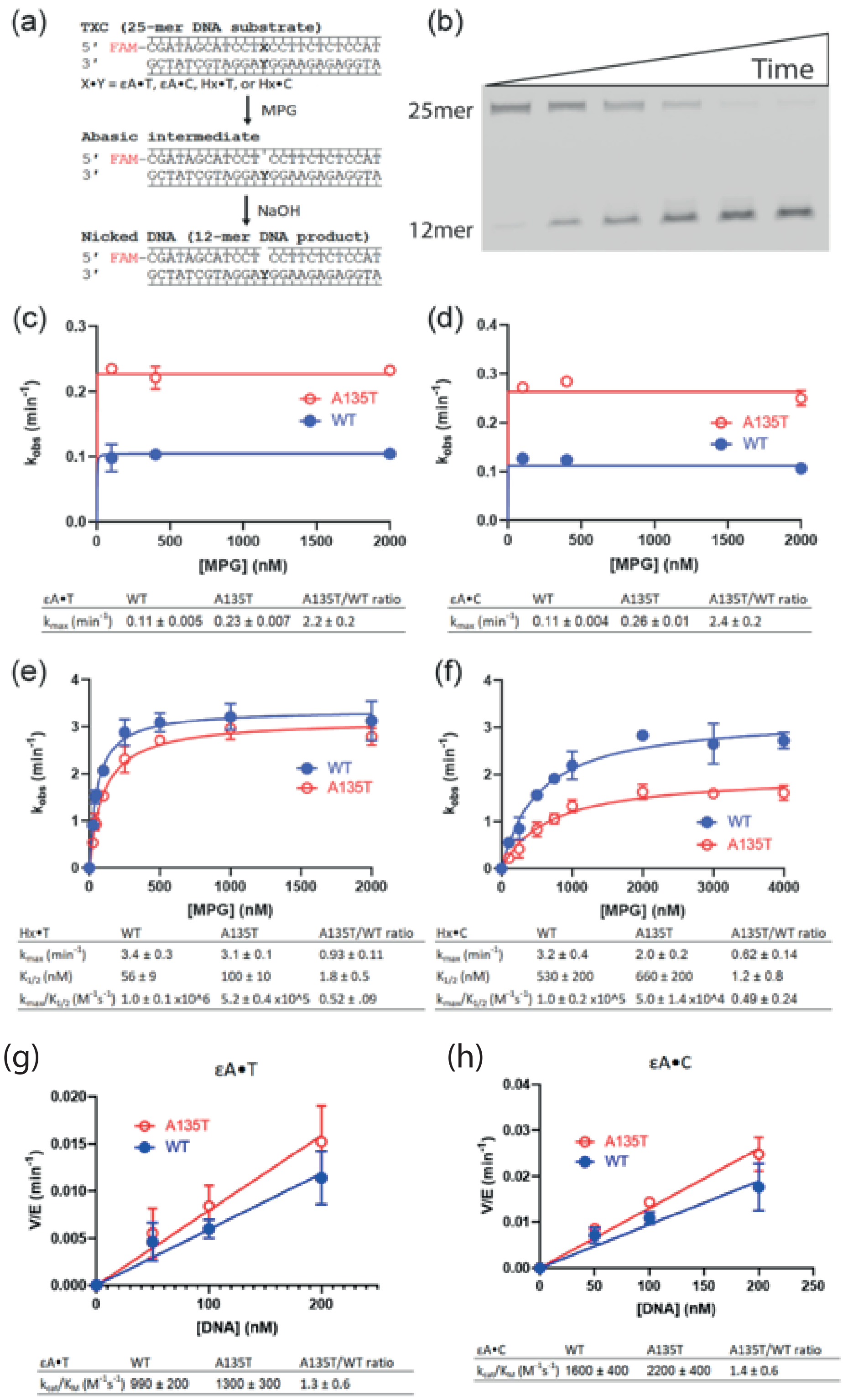


| εA•T | WT | A135T | A135T/WT ratio |
|---|---|---|---|
| $k_{max}$ ($min^{-1}$) | 0.11 ± 0.005 | 0.23 ± 0.007 | 2.2 ± 0.2 |

| εA•C | WT | A135T | A135T/WT ratio |
|---|---|---|---|
| $k_{max}$ ($min^{-1}$) | 0.11 ± 0.004 | 0.26 ± 0.01 | 2.4 ± 0.2 |

| Hx•T | WT | A135T | A135T/WT ratio |
|---|---|---|---|
| $k_{max}$ ($min^{-1}$) | 3.4 ± 0.3 | 3.1 ± 0.1 | 0.93 ± 0.11 |
| $K_{1/2}$ (nM) | 56 ± 9 | 100 ± 10 | 1.8 ± 0.5 |
| $k_{max}/K_{1/2}$ ($M^{-1}s^{-1}$) | 1.0 ± 0.1 x10^6 | 5.2 ± 0.4 x10^5 | 0.52 ± .09 |

| Hx•C | WT | A135T | A135T/WT ratio |
|---|---|---|---|
| $k_{max}$ ($min^{-1}$) | 3.2 ± 0.4 | 2.0 ± 0.2 | 0.82 ± 0.14 |
| $K_{1/2}$ (nM) | 530 ± 200 | 660 ± 200 | 1.2 ± 0.8 |
| $k_{max}/K_{1/2}$ ($M^{-1}s^{-1}$) | 1.0 ± 0.2 x10^5 | 5.0 ± 1.4 x10^4 | 0.49 ± 0.24 |

| εA•T | WT | A135T | A135T/WT ratio |
|---|---|---|---|
| $k_{cat}/K_M$ ($M^{-1}s^{-1}$) | 990 ± 200 | 1300 ± 300 | 1.3 ± 0.6 |

| εA•C | WT | A135T | A135T/WT ratio |
|---|---|---|---|
| $k_{cat}/K_M$ ($M^{-1}s^{-1}$) | 1600 ± 400 | 2200 ± 400 | 1.4 ± 0.6 |

Supplemental Figure 9. In vitro glycosylase activity of WT and A135T MPG. (a) Glycosylase assay for recombinant protein and 25mer lesion-containing oligonucleotides (O'Brien 2003). MPG excises lesion X from X•Y duplex to create an abasic site, which is subsequently hydrolyzed by NaOH to create a 12mer product. (b) Representative denaturing gel scanned for fluorescein fluorescence. (c-d) Concentration dependence for single-turnover excision of

Hx from opposing T and C contexts shows decreased catalytic efficiency for A135T as compared to WT MPG. These single turnover rate constants were fit to the equation kobs = kmax [MPG]/ (K1/2 + [MPG]) as previously described (Lyons 2009). (e-f) Concentration independent excision of εA from opposing T and C shows increased rate of N-glycosidic bond cleavage by A135T. (e-f) Steady state concentration dependence for excision of εA was performed in order to measure the catalytic efficiency (kcat/KM) for A135T and WT MPG using 5 nM enzyme and the indicated concentration of substrate. To circumvent the tight binding by MPG, 800 mM NaCl was added to the standard buffer as previously described, using the equation V/E = kcat/KM[S] (Zhang 2015). Mean ± SD is shown for at least 3 independent experiments.

(a)

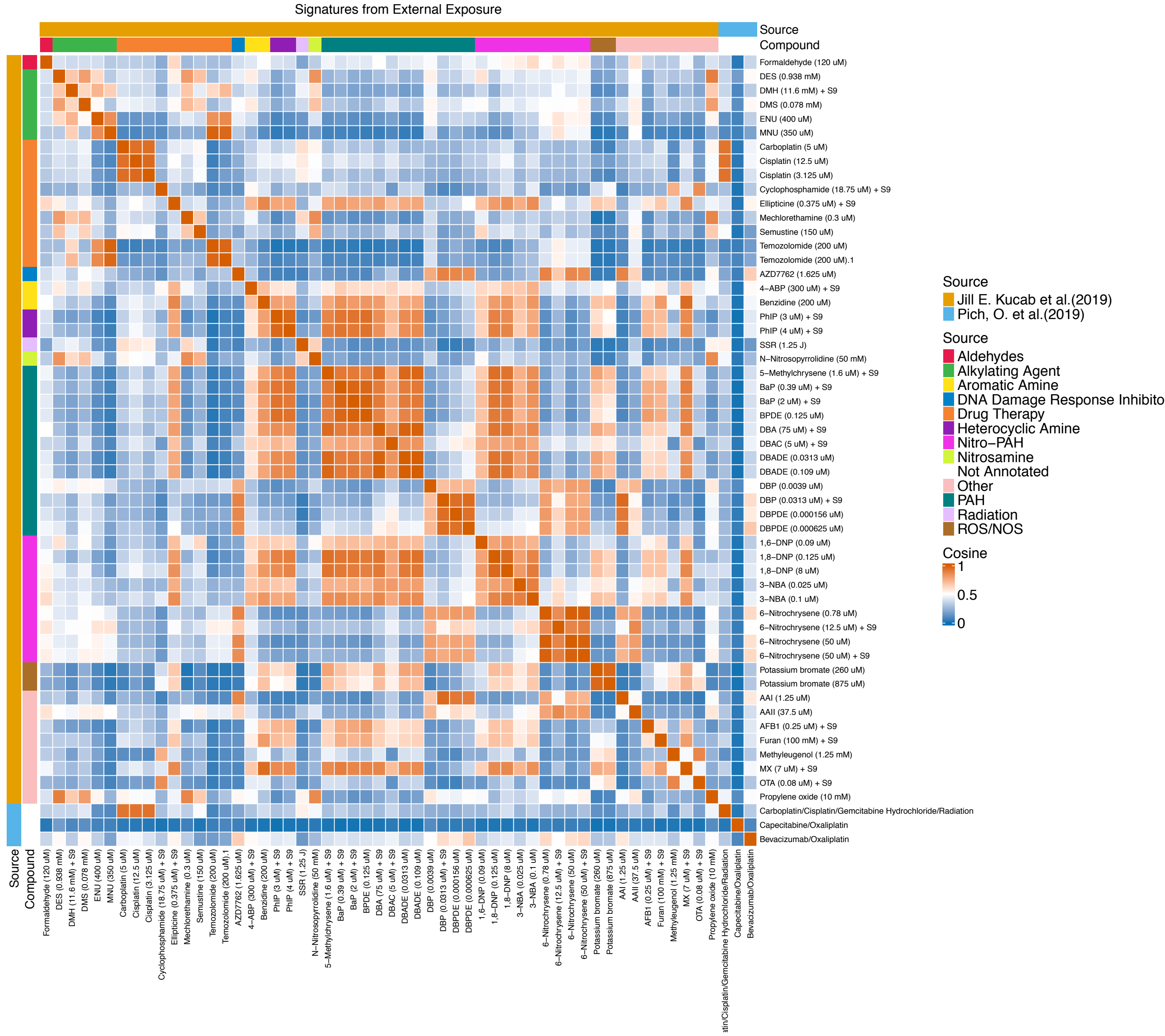
Signatures from External Exposure
Source
Compound
Formaldehyde (120 uM)
DES (0.938 mM)
DMH (11.6 mM) + S9
DMS (0.078 mM)
ENU (400 uM)
MNU (350 uM)
Carboplatin (5 uM)
Cisplatin (12.5 uM)
Cisplatin (3.125 uM)
Cyclophosphamide (18.75 uM) + S9
Ellipticine (0.375 uM) + S9
Mechlorethamine (0.3 uM)
Semustine (150 uM)
Temozolomide (200 uM)
Temozolomide (200 uM).1
AZD7762 (1.625 uM)
4–ABP (300 uM) + S9
Benzidine (200 uM)
PhIP (3 uM) + S9
PhIP (4 uM) + S9
SSR (1.25 J)
N–Nitrosopyrrolidine (50 mM)
5–Methylchrysene (1.6 uM) + S9
BaP (0.39 uM) + S9
BaP (2 uM) + S9
BPDE (0.125 uM)
DBA (75 uM) + S9
DBAC (5 uM) + S9
DBADE (0.0313 uM)
DBADE (0.109 uM)
DBP (0.0039 uM)
DBP (0.0313 uM) + S9
DBPDE (0.000156 uM)
DBPDE (0.000625 uM)
1,6–DNP (0.09 uM)
1,8–DNP (0.125 uM)
1,8–DNP (8 uM)
3–NBA (0.025 uM)
3–NBA (0.1 uM)
6–Nitrochrysene (0.78 uM)
6–Nitrochrysene (12.5 uM) + S9
6–Nitrochrysene (50 uM)
6–Nitrochrysene (50 uM) + S9
Potassium bromate (260 uM)
Potassium bromate (875 uM)
AAI (1.25 uM)
AAII (37.5 uM)
AFB1 (0.25 uM) + S9
Furan (100 mM) + S9
Methyleugenol (1.25 mM)
MX (7 uM) + S9
OTA (0.08 uM) + S9
Propylene oxide (10 mM)
Carboplatin/Cisplatin/Gemcitabine Hydrochloride/Radiation
Capecitabine/Oxaliplatin
Bevacizumab/Oxaliplatin
Source
Jill E. Kucab et al.(2019)
Pich, O. et al.(2019)
Source
Aldehydes
Alkylating Agent
Aromatic Amine
DNA Damage Response Inhibito
Drug Therapy
Heterocyclic Amine
Nitro–PAH
Nitrosamine
Not Annotated
Other
PAH
Radiation
ROS/NOS
Cosine
1
0.5
0

(b)

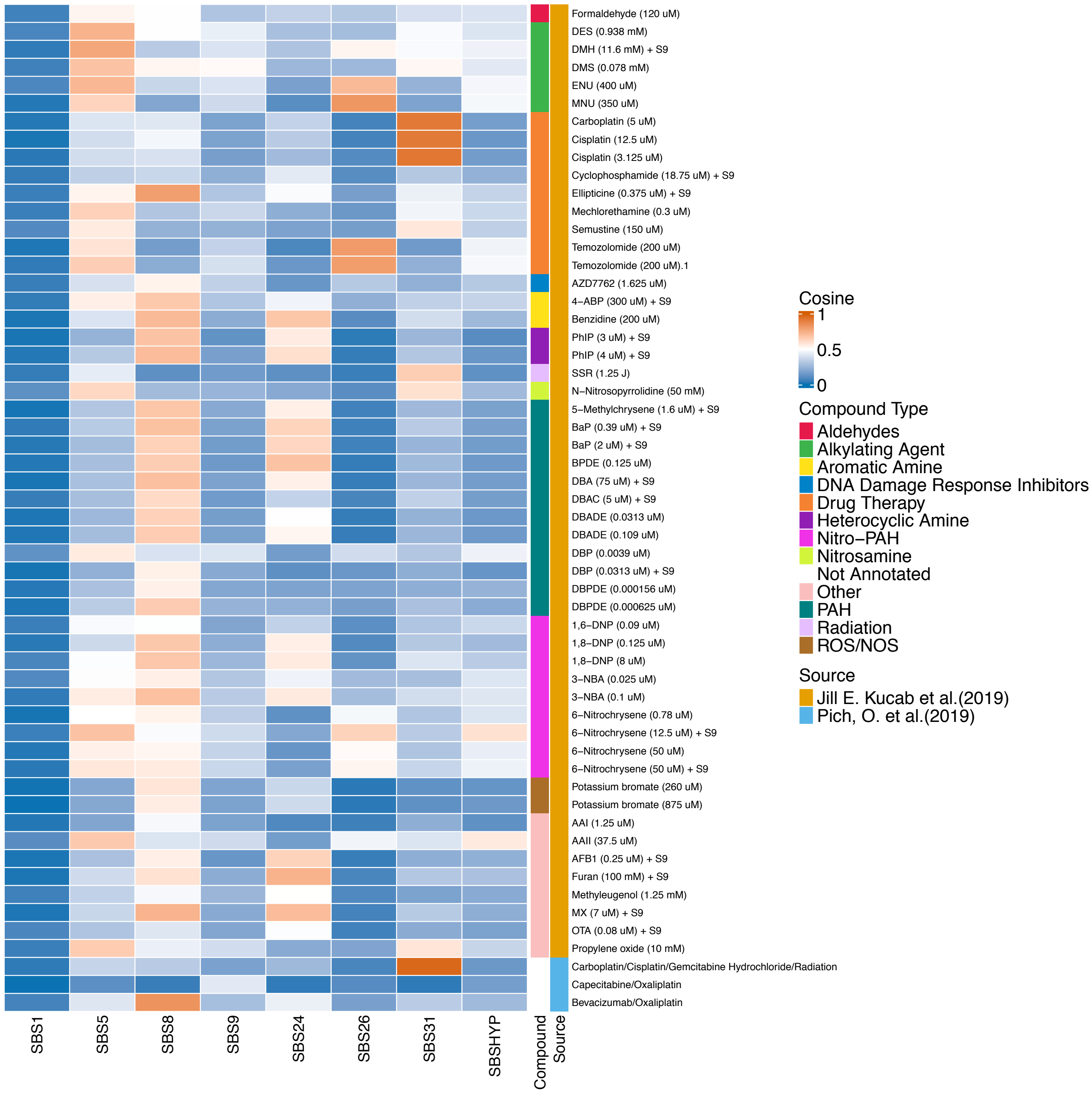


Supplemental Figure 10:

(a) Cosine similarity of all the signatures caused by environmental mutagens amongst themselves.

(b) Cosine similarity of all the signatures caused by environmental mutagens with the extracted signatures from the hypermutated individuals. These signatures were compiled from Kucab et al 2019, Pich et al 2019 and Volkova et al 2020 (see Methods)

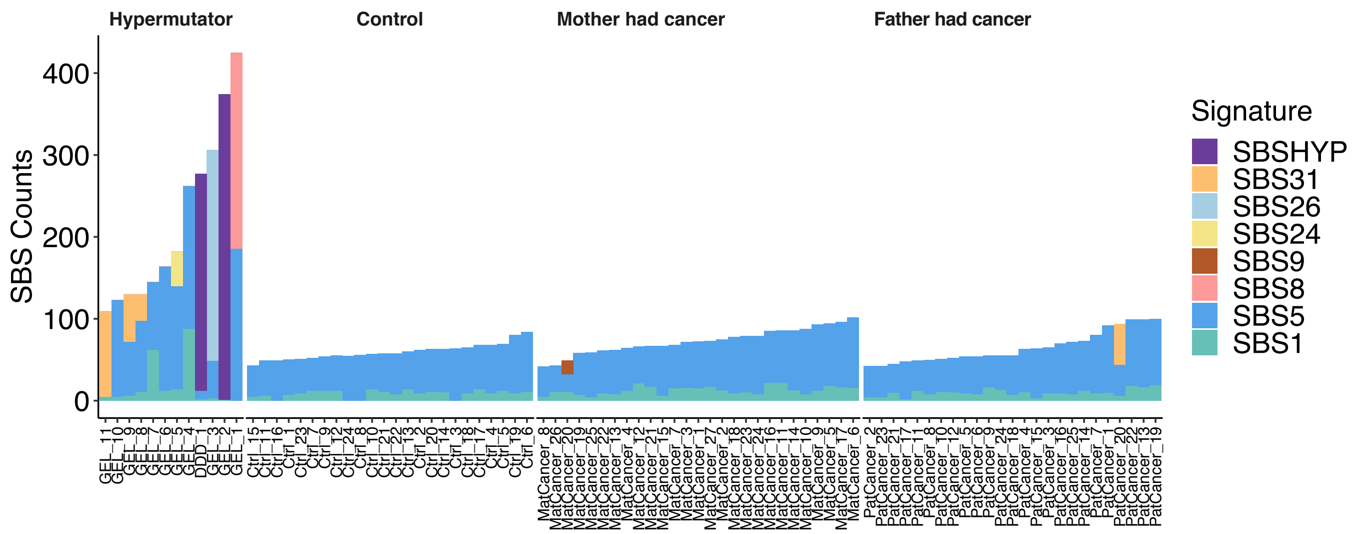


Supplemental Figure 11: Mutational signature contributions for hypermutators, a set of controls selected matched on parental age and individuals who have a parental history of cancer.

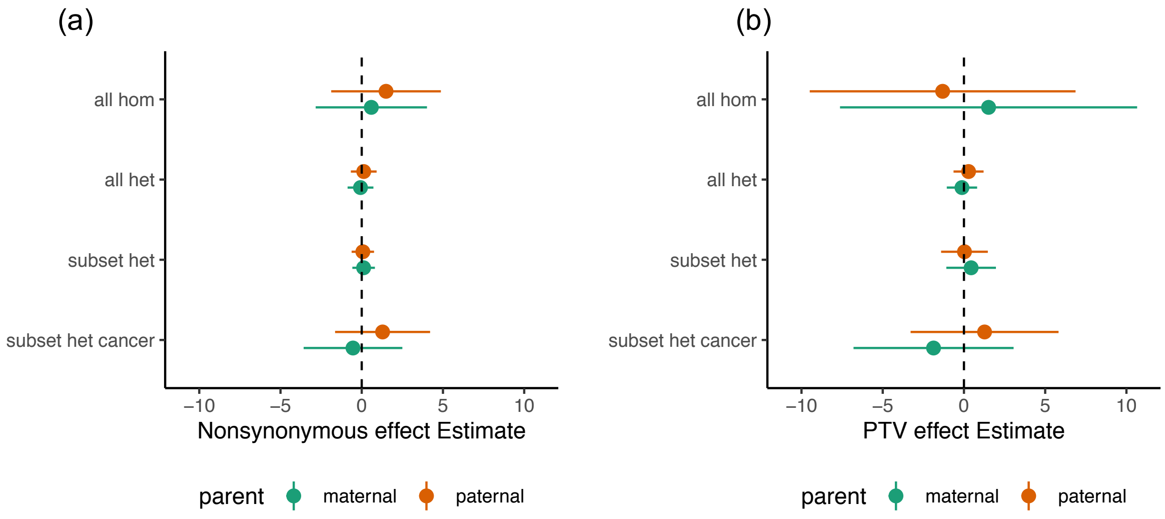


Supplemental Figure 12: Impact of rare variants in DNA repair genes on germline mutation rate. Poisson regression effect estimates for binary variables of having a parental variant in genes known to be involved in DNA repair. (a) considered all nonsynonymous variants in the subsets (b) is restricted to PTVs.

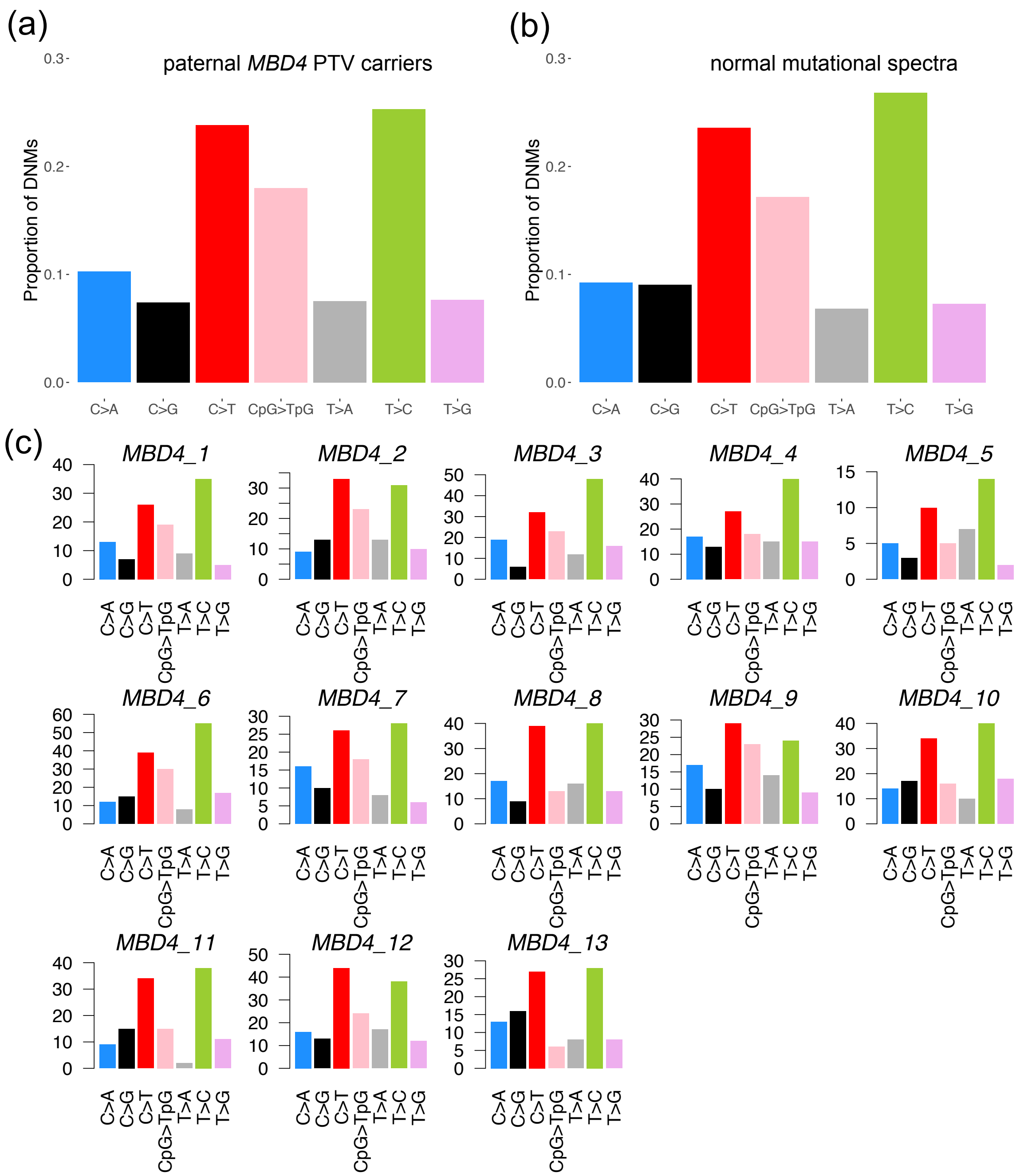


Supplemental Figure 13: Comparing the mutational spectra of DNMs across the 13 paternal *MBD4* paternal PTV carriers (a) with the expected proportion of mutations (b) in each mutation type taken from Rahbari et al. (c) The individual mutational spectra demonstrating that no one individual has an elevated number of CpG>TpG mutations.

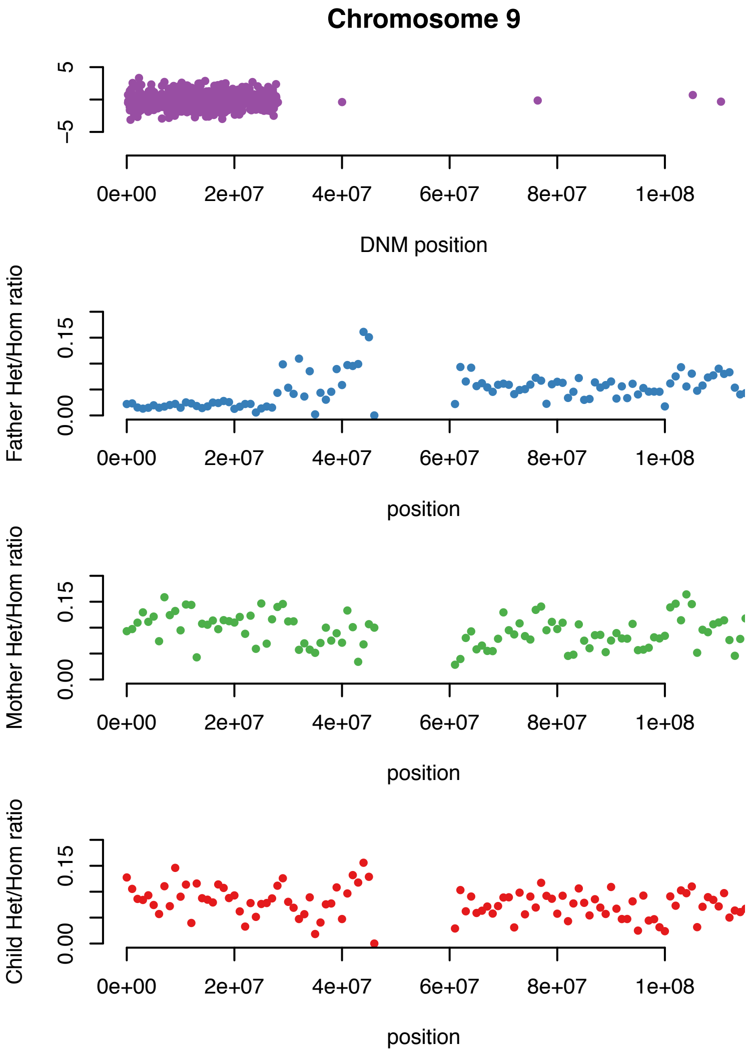


Supplemental Figure 14: Loss of transmitted allele example leading to false positive DNMs Top plot shows the location of the called DNMs in the child on chromosome 9. The plots below show the heterozygous/homozygous ratio in the Father, Mother and Child showing a loss of heterozygosity in the father in the same region the DNMs have been called.

**Supplemental Tables**

Supplemental Table 1: Trinucleotide mutation counts for 12 hypermutated individuals

Supplemental Table 2: Mutation probabilities for novel mutational signature SBSHYP

Supplemental Table 3: DNA repair genes with annotations taken from https://www.mdanderson.org/documents/Labs/Wood-Laboratory/human-dna-repair-genes.html (accessed January 2020)

| ID | Child disease | Genetic variant** | Parental chemotherapy exposure* |
|---|---|---|---|
| GEL_1 | Epileptic encephalopathy | Father: 3:14165549 G>A homozygous<br>NM_004628.5(XPC):c.658C>T (p.Arg220Ter)<br>ClinVar ID: 550020<br>GnomAD allele frequency: 2.2e-5<br>Clinical diagnosis of xeroderma pigmentosum | NA |
| GEL_2 | Multisystem developmental disorder | NA | Nephrotic syndrome: Cyclophosphamide, Chlorambucil (and immunosuppressants) |
| GEL_3 | Intellectual disability | Father: 16:83139 G>A homozygous<br>NM_002434.4(MPG):c.403G>A (p.Ala135Thr)<br>ClinVar ID: absent<br>GnomAD allele frequency: 9.57e-5 | NA |
| GEL_4 | Multisystem developmental disorder, myelodysplasia | Child: 12:11885935 A>G mosaic heterozygous<br>NM_001987.5(ETV6):c.1162A>G (p.Asn388Asp)<br>ClinVar ID: absent<br>GnomAD allele frequency: 0 (absent) | NA |
| GEL_5 | Pulmonary fibrosis | NA | Systemic lupus erythematosus: [Chemotherapy confirmed, drugs unknown] |
| GEL_6 | Congenital myopathy | NA | NA |
| GEL_7 | Intellectual disability | NA | NA |

| | | | |
|---|---|---|---|
| GEL_8 | Abnormality of copper homeostasis | NA | Testicular cancer:<br>Drugs unknown |
| GEL_9 | Intellectual disability | NA | Testicular cancer:<br>BEP (Bleomycin, etoposide and platinum) |
| GEL_10 | Intellectual disability | NA | NA |
| GEL_11 | Cataracts | NA | Cancer of long bones, intestinal tract, lung (secondary):<br>Drugs unknown |
| DDD_1 | Global Developmental Delay, Microcephaly | NA | Hodgkins Lymphoma:<br>ABVD (Bleomycin-Dacarbazine-Doxorubicin-Vinblastine)<br>IVE (Iphosphamide, epirubicin and etoposide) |

*prior to conception
** GRCh38 coordinates

**Supplemental Table 4:** Summary of putative mutagenic variants and parental pharmacological exposures for hypermutated individuals

| MPG variant | alle frequency[1] | eA•T $k_{rel}$ | Hx•T $k_{rel}$ | Specificity eA/Hx | Reference |
|---|---|---|---|---|---|
| R120C[3] | $5{\times}10^{-4}$ | 0.9 | NR[2] | NR | Adhikari, Chetram et al. (2015) |
| Y127W | NR | 0.13 | NR | NR | O'Brien and Ellenberger (2003) |
| R141Q[3] | $8{\times}10^{-5}$ | 0.8 | NR | NR | Adhikari, Chetram et al. (2015) |
| Y159W | NR | 0.37 | NR | NR | O'Brien and Ellenberger (2003) |
| **A135T** | **$1{\times}10^{-4}$** | **2.2** | **0.93** | **2.4** | **this work** |
| R138S | NR | 1.1 | 0.78 | 1.3 | Zhang and O'Brien (2015) |
| R141M | NR | 1.0 | 1.0 | 1.0 | Zhang and O'Brien (2015) |
| R145S | NR | 1.0 | 0.32 | 3.1 | Zhang and O'Brien (2015) |
| Y162W | NR | 1.0 | 0.42 | 2.4 | Hendershot and O'Brien (2017) |
| N169S[4] | NR | 2.0 | 1.1 | 1.8 | O'Brien and Ellenberger (2004) |
| R182M | NR | 0.64 | 0.44 | 1.5 | Zhang and O'Brien (2015) |
| R197S | NR | 1.1 | 0.68 | 1.5 | Zhang and O'Brien (2015) |
| K210M | NR | 0.9 | 1.1 | 0.82 | Zhang and O'Brien (2015) |
| K220M | NR | 1.5 | 0.85 | 1.8 | Zhang and O'Brien (2015) |
| K229M | NR | 1.0 | 0.93 | 1.1 | Zhang and O'Brien (2015) |

**Supplemental Table 5.** Compilation of single turnover excision kinetics for MPG variants. Relative single-turnover glycosylase activity is reported as the ratio of the single turnover rate constant for the variant divided by that of the WT enzyme from the indicated reference. [1]Allele frequency from GnomAD. [2]NR, not reported. [3]R120C and R141Q are the most deleterious variants tested out of 8 rare alleles of MPG. R141Q and to a lesser extent R120C showed a modest increase in mutation frequency in a plasmid repair assay performed in HEK293 cells (Adhikari, Chetram et al. (2015)). [4]N169S shows a mutator phenotype when it is expressed in yeast (Eyler, Burnham et al. (2017), Connor, Wilson et al. (2005)).

| ID | Number SNVs | Number Indels | SNV p-value | Paternal age | Maternal age | Paternal DNMs | Maternal DNMs | Phase p-value | Chemo code |
|---|---|---|---|---|---|---|---|---|---|
| MatCancer_1 | 86 | 5 | 0.014 | (30,35] | (25,30] | 14 | 8 | 0.095 | Y |
| MatCancer_10 | 49 | 4 | 0.983 | (35,40] | (30,35] | 12 | 6 | 0.194 | N |
| MatCancer_11 | 67 | 8 | 0.440 | (30,35] | (30,35] | 15 | 5 | 0.469 | N |
| MatCancer_12 | 61 | 6 | 0.433 | (25,30] | (30,35] | 22 | 3 | 0.940 | N |
| MatCancer_13 | 79 | 10 | 0.249 | (35,40] | (30,35] | 20 | 6 | 0.536 | N |
| MatCancer_14 | 79 | 6 | 0.298 | (35,40] | (35,40] | 15 | 4 | 0.639 | N |
| MatCancer_15 | 60 | 5 | 0.710 | (30,35] | (30,35] | 17 | 5 | 0.562 | N |
| MatCancer_16 | 43 | 6 | 0.609 | (20,25] | (20,25] | 11 | 5 | 0.274 | N |
| MatCancer_17 | 73 | 4 | 0.336 | (35,40] | (30,35] | 19 | 1 | 0.993 | N |
| MatCancer_18 | 72 | 8 | 0.369 | (35,40] | (30,35] | 15 | 7 | 0.201 | N |
| MatCancer_19 | 75 | 11 | 0.318 | (35,40] | (35,40] | 15 | 7 | 0.201 | Y |
| MatCancer_2 | 66 | 4 | 0.464 | (30,35] | (30,35] | 24 | 3 | 0.958 | N |
| MatCancer_20 | 72 | 10 | 0.324 | (30,35] | (30,35] | 18 | 5 | 0.605 | N |
| MatCancer_21 | 64 | 6 | 0.779 | (35,40] | (30,35] | 16 | 2 | 0.934 | N |
| MatCancer_22 | 94 | 6 | 0.155 | (40,45] | (40,45] | 20 | 6 | 0.536 | Y |
| MatCancer_23 | 103 | 4 | 0.101 | (45,50] | (35,40] | 22 | 14 | 0.018 | Y |
| MatCancer_24 | 68 | 6 | 0.553 | (35,40] | (25,30] | 17 | 4 | 0.720 | N |
| MatCancer_25 | 43 | 3 | 0.745 | (20,25] | (20,25] | 11 | 3 | 0.633 | Y |
| MatCancer_26 | 93 | 5 | 0.279 | (45,50] | (30,35] | 22 | 5 | 0.750 | Y |
| MatCancer_27 | 88 | 5 | 0.176 | (40,45] | (30,35] | 17 | 6 | 0.407 | N |
| MatCancer_3 | 62 | 3 | 0.892 | (35,40] | (40,45] | 16 | 3 | 0.829 | Y |
| MatCancer_4 | 86 | 6 | 0.529 | (45,50] | (35,40] | 33 | 8 | 0.721 | N |
| MatCancer_5 | 67 | 5 | 0.565 | (35,40] | (25,30] | 20 | 8 | 0.273 | N |
| MatCancer_6 | 85 | 4 | 0.014 | (25,30] | (30,35] | 21 | 6 | 0.577 | N |
| MatCancer_7 | 96 | 1 | 0.122 | (40,45] | (40,45] | 19 | 10 | 0.091 | N |
| MatCancer_8 | 78 | 3 | 0.092 | (30,35] | (20,25] | 20 | 2 | 0.971 | N |
| MatCancer_9 | 58 | 3 | 0.321 | (25,30] | (20,25] | 11 | 4 | 0.437 | Y |
| PatCancer_1 | 49 | 5 | 0.966 | (30,35] | (30,35] | 12 | 5 | 0.842 | N |
| PatCancer_10 | 94 | 8 | 0.005 | (30,35] | (25,30] | 24 | 3 | 0.118 | N |
| PatCancer_11 | 45 | 10 | 0.713 | (20,25] | (20,25] | 11 | 3 | 0.619 | N |
| PatCancer_12 | 99 | 10 | 0.143 | (45,50] | (25,30] | 21 | 7 | 0.727 | N |
| PatCancer_13 | 42 | 6 | 0.955 | (25,30] | (25,30] | 9 | 6 | 0.968 | N |
| PatCancer_14 | 55 | 7 | 0.800 | (30,35] | (30,35] | 15 | 3 | 0.407 | Y |
| PatCancer_15 | 73 | 10 | 0.147 | (30,35] | (20,25] | 18 | 6 | 0.725 | N |
| PatCancer_16 | 92 | 4 | 0.475 | (55,60] | (25,30] | 27 | 0 | 0.001 | N |
| PatCancer_17 | 42 | 3 | 0.667 | (20,25] | (15,20] | 9 | 4 | 0.858 | N |
| PatCancer_18 | 65 | 3 | 0.806 | (40,45] | (30,35] | 14 | 8 | 0.961 | N |
| PatCancer_19 | 63 | 6 | 0.746 | (35,40] | (30,35] | 14 | 3 | 0.456 | N |
| PatCancer_2 | 52 | 1 | 0.381 | (20,25] | (15,20] | 12 | 3 | 0.563 | Y |
| PatCancer_20 | 54 | 4 | 0.333 | (20,25] | (25,30] | 14 | 7 | 0.925 | N |
| PatCancer_21 | 54 | 4 | 0.828 | (30,35] | (30,35] | 12 | 2 | 0.367 | N |
| PatCancer_22 | 80 | 8 | 0.287 | (35,40] | (30,35] | 16 | 8 | 0.934 | Y |
| PatCancer_23 | 50 | 4 | 0.851 | (25,30] | (30,35] | 12 | 9 | 0.991 | N |
| PatCancer_24 | 55 | 3 | 0.889 | (35,40] | (30,35] | 11 | 7 | 0.970 | Y |
| PatCancer_25 | 51 | 4 | 0.952 | (35,40] | (30,35] | 7 | 5 | 0.968 | N |
| PatCancer_3 | 99 | 6 | 0.129 | (50,55] | (20,25] | 26 | 6 | 0.411 | N |
| PatCancer_4 | 73 | 5 | 0.025 | (25,30] | (20,25] | 20 | 6 | 0.646 | N |
| PatCancer_5 | 64 | 4 | 0.195 | (25,30] | (25,30] | 9 | 3 | 0.732 | N |
| PatCancer_6 | 70 | 5 | 0.721 | (40,45] | (35,40] | 7 | 5 | 0.968 | N |
| PatCancer_7 | 48 | 5 | 0.426 | (20,25] | (15,20] | 17 | 4 | 0.484 | N |
| PatCancer_8 | 55 | 5 | 0.624 | (25,30] | (30,35] | 10 | 6 | 0.954 | N |
| PatCancer_9 | 100 | 5 | 0.001 | (30,35] | (30,35] | 29 | 8 | 0.556 | N |

**Supplementary Table 6:** Individuals with a parent with a cancer diagnosis reported in hospital episode statistics prior to conception. Prefix of ID indicates whether the mother (MatCancer) or father (PatCancer) is the parent with cancer diagnosis. Number of SNVs and Indels refers to the DNM count in the child. The SNV p-value is test for if the number of SNVs is significantly greater than expected given parental age. Paternal and maternal age are given in 5 year bins. The number of paternal and maternal DNMs are the count of DNMs that phased paternally and maternally. The phase p-value is testing if proportion of DNMs that phase paternally is different to overall proportion across 100kGP dataset. Chemo code indicates whether the parent also has an ICD10 code for chemotherapy yes (Y) or no (N).

| Variant Subset | Csq | Genotype | Paternal count | Paternal Effect | Paternal p-value | Maternal count | Maternal effect | Maternal p-value |
|---|---|---|---|---|---|---|---|---|
| All DNA repair | nonsyn | het | 5857 | 0.12 | 0.65 | 5903 | -0.08 | 0.79 |
| | PTV | het | 1203 | 0.28 | 0.36 | 1150 | -0.12 | 0.70 |
| | nonsyn | hom | 78 | 1.50 | 0.19 | 71 | 0.59 | 0.61 |
| | PTV | hom | 13 | -1.31 | 0.64 | 11 | 1.52 | 0.62 |
| Subset DNA repair | nonsyn | het | 3075 | 0.07 | 0.77 | 2918 | 0.12 | 0.62 |
| | PTV | het | 432 | 0.03 | 0.95 | 388 | 0.44 | 0.39 |
| Germline cancer | nonsyn | het | 103 | 1.28 | 0.19 | 97 | -0.54 | 0.60 |
| | PTV | het | 41 | 1.27 | 0.41 | 35 | -1.87 | 0.26 |

**Supplemental Table 7**: Impact of parental rare variants in DNA repair genes on germline mutation rate. Effect estimates and corresponding p-values from 8 regression models on three subsets of variant groups. Csq: consequence of variants examined where ‘nonsyn’ refers to nonsynonymous variants and PTV refers to a subset of these of just protein truncating variants. Genotype details whether the variants considered were ‘het’: heterozygous or ‘hom’: homozygous. Paternal count refers to the number of variants found in this subset in paternal genomes and maternal count refers to the equivalent for mothers.

| MAF bin | LD group | Maternal $h^2$ | Maternal SE | Paternal $h^2$ | Paternal SE |
|---|---|---|---|---|---|
| 0.001-0.01 | low | 0.151 | 0.195 | 0.337 | 0.167 |
| | High | -0.008 | 0.205 | 0.181 | 0.136 |
| 0.01-0.05 | low | -0.018 | 0.083 | $10^{-6}$ | 0.070 |
| | high | 0.026 | 0.032 | $10^{-6}$ | 0.026 |
| 0.05-1 | low | -0.074 | 0.061 | $10^{-6}$ | 0.051 |
| | high | -0.002 | 0.032 | 0.008 | 0.027 |
| TOTAL | - | 0.071<br>p = 0.21 | 0.255 | 0.526<br>p = 0.095 | 0.165 |
| Number of individuals | - | 6329 | | 6352 | |

**Supplemental Table 8:** Maternal and paternal SNP heritability of residuals of number of dnSNVs after correcting for parental age, hypermutation status and data quality. Results from GREML-LDMS binned on three minor allele frequency (MAF) bins and two LD groups. High LD refers to variants with LD > median LD and low LD refers to variants with LD < medianLD. Maternal heritability has negative estimates as this was run without being constrained to positive numbers due to estimates not converging otherwise. SE refers to standard error of the $h^2$ estimate. Performed on a subset of individuals with white british ancestry.
